# GlycoVHH: Introducing N-glycans on the camelid VHH antibody scaffold - Optimal sites and use for macrophage delivery

**DOI:** 10.1101/2022.04.25.489312

**Authors:** Loes van Schie, Wander Van Breedam, Charlotte Roels, Bert Schepens, Martin Frank, Ahmad Reza Mehdipour, Bram Laukens, Wim Nerinckx, Francis Santens, Simon Devos, Iebe Rossey, Karel Thooft, Sandrine Vanmarcke, Annelies Van Hecke, Nico Callewaert

**Affiliations:** UGent Department of Biochemistry and microbiology, Ghent University, Ghent, Belgium; VIB-UGent Center for Medical Biotechnology, Ghent, Belgium; Biognos AB, Göteborg, Sweden; UGent Center for Molecular Modeling, Ghent University, Ghent, Belgium; UGent Department of Organic and Macromolecular Chemistry, Ghent University

**Keywords:** VHHs, N-glycosylation, *Pichia pastoris* GlycoSwitchM5, macrophage targeting, single-domain antibody scaffold

## Abstract

As small and stable high-affinity antigen binders, VHHs boast attractive characteristics both for therapeutic use in various disease indications, and as versatile reagents in research and diagnostics. To further increase the versatility of VHHs, we explored the VHH scaffold in a structure-guided approach to select regions where the introduction of an N-glycosylation N-X-T sequon and its associated glycan should not interfere with protein folding or epitope recognition. We expressed variants of such glycoengineered VHHs in the *Pichia pastoris* GlycoSwitchM5 strain, allowing us to pinpoint preferred sites at which Man_5_GlcNAc_2_-glycans can be introduced at high site occupancy without affecting antigen binding. A VHH carrying predominantly a Man_5_GlcNAc_2_ N-glycan at one of these preferred sites showed highly efficient, glycan-dependent uptake by Mf4/4 macrophages *in vitro* and by alveolar lung macrophages *in vivo,* illustrating one potential application of glyco-engineered VHHs: a glycan-based targeting approach for lung macrophage endolysosomal system delivery. The set of optimal artificial VHH N-glycosylation sites identified in this study can serve as a blueprint for targeted glyco-engineering of other VHHs, enabling site-specific functionalization through the rapidly expanding toolbox of synthetic glycobiology.

## Introduction

Since the 1980s, recombinant antibodies have gradually acquired a dominant position in the fields of biotherapeutics and biotechnological detection and imaging. In the classic antibody, antigen binding capacity is spread over the N-terminal variable domains of both heavy and light chains. In the early 1990s however, + discovered functional antibodies in camelid species that lack a light chain. Intriguingly, the single N-terminal domains of these antibodies turned out to be both stable and sufficient for antigen binding without requiring pairing to other domains, and they were rapidly adopted as next generation immune reagents, designated as VHHs (for the Variable domain of the Heavy chain of Heavy chain-only antibodies, also known as single-chain antibodies or Nanobodies^®^). As in regular antibody domains, the beta-sandwich scaffold structure is generally conserved in these VHHs, whereas the complementarity-determining regions (CDRs) that bind antigen are highly variable to allow specific binding to a wide range of targets (Desmyter et al., 1996).

The VHH antibody format boasts a set of attractive characteristics, including small size, high physicochemical stability, manufacturability in microorganisms, and limited immunogenicity upon straightforward humanization, making them suited for a wide range of biotechnological and therapeutic applications. In nature, only a minority of VHHs contain N-glycans, mostly when selected by chance as part of the antigen-binding CDRs (Harmsen et al., 2009). This (CDR) glycosylation is not preferred in terms of easy developability of the VHH, as glycan structures are often heterogeneous and different across expression systems, leading to heterogeneous antigen binding behavior. However, the introduction of N-glycans on the scaffold rather than in the CDR could enable glycan structure synthetic biology and glycan-targeted coupling chemistry according to emerging approaches (Baskin and Bertozzi, 2007; Jacobs et al., 2009; Meuris et al., 2014; Van Breedam et al., 2021) to import novel functionality to VHHs, without affecting antigen binding.

For example, biotherapeutics sometimes need to be endocytosed by specific cells to be effective, which can be achieved by glycan-targeting of the therapeutic protein to cell-type specific endocytic receptors. A noteworthy example in this context is the macrophage mannose receptor (MMR), a macrophage-expressed mannose-binding lectin. In Gaucher disease, the glucocerebrosidase storage product is mainly located in macrophage lysosomes, and therapy with mannosylated MMR-targeting enzyme replacement therapy is effective (Shaaltiel et al., 2007; Zimran et al., 2018). Also, pathogens such as *Mycobacterium, Listeria, Salmonella, Legionella, Brucella* and *Acetinobacter baumanni* can infect macrophages and survive and propagate intracellularly, within the phagolysosomal compartment (Sycz et al., 2021). Hence, technology for targeting of antimicrobial agents to the macrophage phagolysosomal compartment is important. Antibiotics have been targeted to the MMR by incorporation into mannosylated carriers such as liposomes, microparticles, nanoparticles or dendrimers (Azad et al., 2014; Kumar et al., 2006; Vieira et al., 2018). An expansion of this concept to VHHs could enable new antibody-mediated antimicrobial agents (Lehar et al., 2015).

Considering the advent of VHHs in therapy (Scully et al., 2019) and inspired by the concept of lectin receptor-directed targeting using mannose-terminal N-glycans, we aimed to introduce high-mannose glycans on the conserved VHH scaffold. In an extensive structure-guided approach, we selected loop regions in the VHH scaffold where the introduction of an N-linked glycan had a chance of not interfering with protein folding or epitope recognition. Selected sites were introduced in an anti-GFP benchmark VHH protein (GFP-binding protein, GBP), upon which quantitative glycosylation site-occupancy and effects on protein stability and antigen recognition were evaluated. Several of the best sites were validated in more biomedically relevant VHHs, and we demonstrated glycan-dependent uptake by macrophages both *in vitro* and *in vivo* in the lung.

## Materials and methods

Throughout the manuscript, the AHo scheme (Honegger and Plückthun, 2001) is used to number immunoglobulin domain amino acid residues.

### Construct design, protein production and protein purification

Plasmids encoding a first set of GBP glycovariants were constructed via Gibson assembly: cDNAs encoding the GBP variants were ordered as synthetic DNA fragments (IDT gBlocks) and cloned into the pKai61 vector (Schoonooghe et al., 2009; sequences in Supplementary A). Plasmids encoding additional GBP glycovariants, F-VHH-4 and F-VHH-L66 glycovariants (Rossey et al., 2017), and AA6 glycovariants (Sequence ID 5 in WO 2017/066468; Feng et al., 2017) were created by introducing synthetic DNA fragments (IDT gBlocks and BioXP DNA tiles) into a Golden Gate assembly-based modular cloning system (Lee et al., 2015). As a result of the modular cloning strategy used and incomplete processing of the N-terminal *S. cerevisiae* α-mating factor secretion signal, a variable (Glu-Ala)-Glu-Ala-Gly-Ser sequence remained at the N-terminus of the second set of GBP glycovariants. All cloned VHHs have a C-terminal His6 (first set of GBP glycovariants) or His8 (all other VHHs) tag and are under control of the methanol-inducible AOX1 promoter and an N-terminal *S. cerevisiae* α-mating factor or oligosaccharyl transferase (Ost1) secretion signal sequence. Plasmids encoding VHH glycovariants were transformed into the GlycoSwitchM5 strain (Jacobs et al., 2009) of *P. pastoris* (re-classified as *Komagataella phaffi;* Kurtzman, 2009). Transformed *P. pastoris* cultures were propagated for 48h at 28 °C in BMGY medium (100 mM potassium phosphate pH 6, 2%_w/v_ peptone, 1%_w/v_ yeast extract, 1%_w/v_ yeast nitrogen base without amino acids and 1% glycerol) containing glycerol as the sole carbon source, and subsequently transferred to BMMY medium (100 mM potassium phosphate pH 6, 2%_w/v_ peptone, 1%_w/v_ yeast extract, 1%_w/v_ yeast nitrogen base without amino acids and 1% methanol) to induce the AOX1 promoter. After 48 hours of incubation at 28 °C, during which 1% methanol was spiked twice daily, supernatant of each recombinant culture was collected. The supernatants were either analyzed directly, or VHHs were purified via nickel metal affinity chromatography (IMAC) over a HisTrap colum (GE Healthcare) and size exclusion chromatography (SEC) over a Superdex 75 column (GE Healthcare) prior to further analysis. Purified proteins were aliquoted, snap-frozen in liquid nitrogen, and stored at −80 °C.

A construct encoding *A. victoria* GFP (UniProt P42212 with Q80R mutation) flanked N-terminally by a His6 tag and an AviTag™ peptide (GLNDIFEAQKIEWHE; Fairhead & Howarth, 2015) was cloned into the pLH36 vector for bacterial protein expression. The GFP-encoding plasmid was co-transformed with the pBirAcm plasmid encoding BirA biotin ligase (Avidity LLC, Colorado) into BL21(DE3) heat-shock competent *E. coli* cells. *E. coli* was propagated in terrific broth (Sigma) at 37 °C until OD_600_ 0.5, after which recombinant protein expression was induced by addition of 0.5 mM IPTG (Sigma) and shaking incubation for 20 more hours at 28 °C or 37 °C. Cell pellets were harvested by centrifugation and lysed by sonication (30 s on, 30 s off for 3.5 minutes at 70% amplitude in a Q500 sonicator, QSonica) in lysis buffer (400 mM NaCl, 20 mM Na_2_HPO_4_/NaH_2_PO_4_ pH 7.5, 20 mM imidazole, 5 mM MgCl_2_, 0.1% Triton X-100 and 1X Roche c0mplete™ Protease Inhibitor). The lysate was cleared by 15 minutes of centrifugation at 48,000×*g*. GFP was purified from the cleared lysate via IMAC (HisTrap) and SEC (Superdex 75), snap-frozen in liquid nitrogen, and stored at −80 °C.

### Glyco-VHH characterization

To assess the presence of N-linked glycans, samples of *P. pastoris* supernatant or purified protein were heat-denatured in buffer containing 5% SDS and 400 mM DTT, treated with PNGaseF, *H. jecorina* endoT (Stals et al., 2010), endoH (New England Biolabs) or mock treated, and assayed via Coomassie Brilliant Blue-stained SDS-PAGE and His-tag-specific western blot. Selected glycovariants were further characterized by intact protein mass spectrometry. LC-MS was performed on an Ultimate 3000 HPLC (Thermo Fisher Scientific, Bremen, Germany) equipped with a Poroshell 300SB-C8 column (Thermo Scientific 1.0 mm of I.D. × 150 mm), in-line connected with an ESI source to an LTQ XL mass spectrometer (Thermo Fischer Scientific). Mobile phases were 0.1% formic acid and 0.05% trifluoroacetic acid (TFA) in H_2_O (solvent A) and 0.1% formic acid and 0.05% TFA in acetonitrile (solvent B). The proteins (as indicated in the Results section: in 10 μl of spent *P. pastoris* supernatant or purified at 1 mg/ml) were separated using a gradient ranging from 10% to 80% solvent B at a flow rate of 100 μl/min for 15 min. The mass spectrometer was operated in MS1 profile mode in the orbitrap analyzer at a resolution of 60,000 (at m/z 400) and a mass range of 600-4000 m/z. The following ESI parameters were used: a surface-induced dissociation of 30 V, a spray voltage of 5.0 kV, capillary temperature of 325 °C, capillary voltage of 35 V and a sheath gas (N2) flow rate setting of 7.

To evaluate the thermal stability of the different glycovariants, differential scanning fluorimetry was performed (Lo et al., 2004). Briefly, a 10 μM solution of GBP-WT or selected glycovariants in HEPES-buffered saline (HBS, 20 mM HEPES pH 7 and 150 mM NaCl) was mixed with 10X SYPRO Orange dye (Life Technologies), and dye binding to protein was measured over a temperature gradient from 20 °C to 98 °C in a Roche LightCycler 480 qPCR machine, after which averages of triplicate melting curves were blank-subtracted and each normalized to 100% fluorescence. T_m_-values were determined as the temperature at 50% fluorescence in the ascending part of the melting curve.

The GFP-binding characteristics of GBP glycovariants were assessed via biolayer interferometry. Briefly, 100 nM of biotinylated Avitag®-GFP was immobilized on a streptavidin-coated biosensor (ForteBio). Two-fold dilution series (8 to 0.125 nM) of the GBP glycovariants were prepared in 20 mM HEPES pH 7, 150 mM NaCl, 0.01% bovine serum albumin and 0.002% Tween-20, and affinity for GFP was measured at 25 °C. Biosensors were regenerated by three times 5 s exposure to 0.1 M glycine pH 3. Data were analyzed using ForteBio Data Analysis 9.0 software. Reported K_d_, k_on_ and k_off_ values are the averages and standard deviations of two or three replicate experiments.

Glycosylation analysis by capillary electrophoresis (DSA-FACE) was carried out as described by Laroy et al. (2006). Briefly, glycoproteins were immobilized on 96 well-plates containing a hydrophobic immobilon-P PVDF membrane (Merck) and their glycans were released by PNGaseF treatment (in-house produced). These N-glycans were derivatized with an APTS label (10 mM) by reductive amination and excess label was removed by SEC over Sephadex G10 resin (GE Healthcare). Labeled glycans were analyzed on an ABI3130 Genetic Analyzer (Applied Biosystems). As internal references, partially hydrolyzed *Leuconostoc mesenteroides* dextran and N-glycans from bovine pancreas RnaseB were included. Exoglycosidase digests were performed on labeled glycans overnight in 5 mM NH_4_OAc pH 5.2, with *H. jecorina* α-1,2-mannosidase (in-house produced, 0.33 μg per digest) and Jack Bean α-1,2/3/6-mannosidase (in-house produced, 19 mU per digest).

To assess neutralization of respiratory syncytial virus (RSV), dilution series of F-VHH-4 and F-VHH-L66 were prepared in Opti-MEM (Gibco), incubated with RSV strain A2 for 30 minutes at 37 °C, and used to infect confluent Vero cells. After three hours, an equal volume of 1.2% Avicel RC-581 (FMC Biopolymers) in DMEM medium supplemented with 2% FCS, 2 mM l-glutamine, non-essential amino acids and 1 mM sodium pyruvate was added to each well and the infection was allowed to continue at 37 °C for three days. Viral infection was tested by immunostaining of the viral plaques with goat anti-RSV serum (AB1128, Chemicon International) and horseradish peroxidase-conjugated anti-goat IgG (SC2020, Santa Cruz). The plaques were visualized by applying TrueBlue peroxidase substrate (KPL, Gaithersburg).

### Molecular modeling

To assess the dynamics of N-glycosylated VHHs, molecular dynamics simulations were performed. To first assess the effect of the glycans on protein surface accessibility, we used the GlycoSHIELD approach (Gecht et al., 2021) for both GlycoSwitchM5 Man_5_GlcNAc_2_ and yeast Golgi-type Man_9_GlcNAc_2_ glycans. Since the Man_9_GlcNAc_2_ glycan was not present in the glycan library of GlycoSHIELD, we added its conformational ensemble to the library. For the addition, we grafted the Man_9_GlcNAc_2_ glycan to a Gly-Asn-Gly tripeptide and then ran a simulation of 3.4 μs in a solvent box using Gromacs v2020.2 (Abraham et al., 2015). Then, an ensemble of glycan conformations was generated based on extracting snapshots from the trajectory at 100 ps intervals.

Later, both glycans’ conformation libraries were grafted into each of the glycosylation sites Q14N, G27N, P48N, R86N, K97N and E99N (AHo numbering) in a modified structure of GBP (PDB id 3OGO, chain E; Kubala et al., 2010). These sites correspond to Q14N, G27N, P42N, R77N, K88N and E90N in the differently numbered PDB-structure. The solvent accessible surface area (SASA) of protein surface was calculated using the GlycoSHIELD script for four different probe sizes of 1.4, 2.5, 7.5, and 10.0 Å. Finally, the shielding of the protein surface by glycans at each glycosylation site was calculated using the following equation:

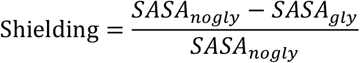

in which SASA_nogly_ and SASA_gly_ are the SASA of the protein surface for the unglycosylated and glycosylated protein, respectively.

To more thoroughly assess the space occupancy at each of the N-glycosylation sites, the structure of GBP was modified using YASARA (Krieger and Vriend, 2014) to include single glycosites Q14N-P15A-G16T, G27N-P30T, P48N-K50T, R86N, K97N-P98A-E99T and E99N (AHo numbering), respectively. These sites correspond to Q14N-P15A-G16T, G27N-P30T, P42N-K54T, R77N, K88N-P89A-E90T and E90N in the differently numbered PDB-structure. A yeast Golgi-type Man_9_GlcNAc_2_ glycan was attached to the newly created single glycosites in each of the six GBP variants (N14, N27, N42, N86, N97 and N99, respectively). The glycoprotein structures were solvated in a box of water with 0.9% NaCl. Molecular dynamics simulations were performed using periodic boundary conditions at 310 K with YASARA (Krieger and Vriend, 2015) in ‘fast-mode’ using high-performance multicore CPUs with GPU support. The AMBER14 force field (Maier et al., 2015) was used for the protein and GLYCAM06 (Kirschner et al., 2008) for the glycan. Two trajectories of 1 μs were sampled for each of the six molecular systems. The space occupancy of the glycans was investigated based on trajectories covering a time scale of 2 μs for each system using Conformational Analysis Tools (www.md-simulations.de/CAT/) in order to understand better which regions of the protein surface are modelled to be occluded by the glycans.

### Uptake of GBP glycovariants by alveolar macrophages in vitro

GBP-WT, GBP-G27N-P30T and GBP-R86N (150 μM in HBS pH 7.5) were fluorescently labeled with an 8-fold molar excess of NHS-modified Alexa Fluor 488 (AF488) dye (Molecular Probes). Labeling mixtures were incubated for 1 hour at room temperature while shaking and subsequently kept at 4 °C until SEC (Superdex 75, equilibrated in HBS pH 7.5). The degree of labeling (DOL) was calculated according to manufacturer’s instructions using the absorbance at 280 and 494 nm and labeling was confirmed by a titration of the labeled His-tagged GBP variants on a nickel-NTA coated plate, with fluorescence detection. Labeled proteins were aliquoted, snap-frozen in liquid nitrogen, and stored at −70 °C.

HEK293T cells were cultivated at 37°C in the presence of 5% CO_2_ in DMEM supplemented with 10% heat-inactivated fetal calf serum (FCS), 1% penicillin, 1% streptomycin, 2 mM L-glutamine, non-essential amino acids (Invitrogen, Carlsbad, California), and 1 mM sodium pyruvate. Mf4/4 cells were grown at 37°C in the presence of 5% CO_2_ in RPMI supplemented with 10% heat-inactivated fetal calf serum (FCS), 1% penicillin, 1% streptomycin, 2 mM L-glutamine, non-essential amino acids (Invitrogen, Carlsbad, California), 1 mM sodium pyruvate and 20 mM HEPES (Gibco).

For a flow cytometry GBP uptake assay, a mixture of 50,000 HEK293T cells and 40,000 Mf4/4 cells was seeded in a 12-well plate in Mf4/4 growth medium. Forty-eight hours after cell seeding, PBS, GBP-WT or GBP-G27N-P30T (0.125-12.5 μg/ml) was added to the co-cultures. After 40 minutes at 37°C, cells were washed twice with ice-cold PBS and detached with trypsin at room temperature. After washing twice with ice-cold PBS, aspecific protein binding capacity of the cells was blocked with 1% BSA, and cells were stained with an AF647 conjugated anti-CD45 antibody (BioLegend, San Diego, California) and analyzed on an BD LSRII flow cytometer. Data analysis was performed using Flowing Software 2.5.1.

For microscopic analysis, Mf4/4 cells were seeded in a μ-slide 8-well (ibidi) and labeled with 100 nM LysoTracker Red (ThermoFisher Scientific). Two hours later the cells were incubated with PBS or 12.5 μg/ml GBP-WT, GBP-G27N-P30T or GBP-R86N for 40 minutes at 37 °C. After washing twice with PBS, the cells were fixed with 1% PFA for 20 minutes, washed twice with PBS, blocked with 1% BSA, and stained with an AF647 conjugated anti-CD45 antibody (BioLegend, San Diego, California). The nuclei were counterstained with Hoechst. The samples were imaged on a LSM880 Airyscan (Carl Zeiss, Jena, Germany) with a 63× PlanApo NA:1.4, oil immersion, DIC M27. The operating software was ZEN blue 2.3. 25 Z-slices were taken, each 0.16 μm, with the Airyscan detector in the super-resolution mode (SR) resulting in a voxel size of 0.04 × 0.04 × 0.16 μm. A 3D deconvolution step with a Wiener filter was carried out post-acquisition using ZEN Blue software.

For live cell imaging, Mf4/4 macrophages were seeded at 50,000 cells per well in ibidi Flow Chambers μ-Slides VI in Mf4/4 growth medium. Cells were incubated with LysoTracker for 30 min and washed three times with culture medium. The flow chamber was mounted on an observer Z. 1 microscope equipped with a Yokogawa disk CSU-X1 (Zeiss, Belgium) equipped with an incubation chamber at 37 °C with 5% CO_2_.

Fluorescence and DIC images were captured using a pln Apo 40x/1.4 oil DIC III objective and a Rolera em-c_2_ Camera. Two selected fields of cells per condition were imaged every 5 min for a total period of 1 hour.

After 1 min of imaging, 12.5 μg/ml of Alexa Fluor 488-labeled GBP-WT, GBP-G27N-P30T or GBP-R86N was administered to the cells via perfusion at a rate of 1 ml/min for 1 min. After 20 min of incubation, VHHs were washed out via perfusion of culture medium at a rate of 0.1 ml/min.

### Uptake of GBP glycovariants by alveolar macrophages in vivo

All animal experiments described in this study were conducted according to national legislation (Belgian laws 14/08/1986 and 22/12/2003, Belgian Royal Decree 06/04/2010) and European legislation (EU Directives 2010/63/EU and 86/609/EEC). All experiments on mice and animal protocols were approved by the ethics committee of Ghent University (permit numbers LA1400091 and EC2018-49). Specific pathogen-free (SPF) female BALB/c mice of 7–8 weeks were purchased from Charles River and housed in a biosafety-level-2, SPF temperature-controlled environment with 12 h light/dark cycles and water and food *ad libitum*. After adaptation in the animal room for 3 weeks, mice were anesthetized with isoflurane (Abbott Animal Health) and treated intranasally with PBS or 12.5 μg of either GBP-WT or GBP-G27N-P30T. One hour after administration, bronchoalveolar lavage was performed. Mice were terminally sedated via intraperitoneal injection of Nembutal (pentobarbital; 125 mg/kg in PBS; Lundbeck, Valby, Denmark). A small incision was made in the trachea to put a lavage cannula in the trachea. Lungs were lavaged 3 times with 1 ml of HBSS with 0.05 mM EDTA (Sigma-Aldrich) and the bronchoalveolar lavage fluid (BALF) was kept on ice. After washing once with ice cold PBS, the majority (75%) of the cells present in the bronchoalveolar lavage fluid (BALF) were stained with either an APC-conjugated anti-mouse F4/80 antibody (eBioscience) in combination with a V450 conjugated anti-mouse CD45 antibody (BD Biosciences) or alternatively with an AF647-conjugated anti-mouse CD45 antibody (Biolegend) in PBS containing 1% BSA (PBS/BSA) for 1 hour. Subsequently, the cells were washed 3 times with PBS/BSA and analyzed on an BD LSRII flow cytometer. All staining procedures were performed using ice-cold buffers. Data analysis was performed using Flowing Software 2.5.1. The remaining BALF cells were allowed to attach to the bottom of a μ-slide 8-well in Mf4/4 growth medium by sequential incubation at 4 °C and 37 °C for 1 hour. Subsequently, the cells were washed once with PBS, fixed with 2% PFA for 20 minutes and washed 3 times with PBS. After blocking for 1 hour with PBS/BSA, the cells were stained with Hoechst nuclear dye, AF647 conjugated anti-mouse CD45 and biotin conjugated anti-mouse F4/80 (Thermo Fisher Scientific, Waltham, MA, USA). The binding of anti-mouse F4/80 was detected using AF568 conjugated streptavidin (Thermo Fisher Scientific, Waltham, MA, USA). Finally, the samples were washed twice with PBS/BSA and 3 times with PBS. Confocal images were taken using a Leica SP5 confocal microscope (Leica Microsystems, Germany).

## Results

### Rational design and production of N-glycosylated VHHs in the yeast strain P. pastoris GlycoSwitchM5

In order to find regions suitable for the introduction of glycans, we studied the structure of a benchmark VHH (GFP-binding protein, GBP; Kubala et al., 2010). Avoiding the beta sandwich core that provides stability to the immunoglobulin fold, we identified the VHH regions that we assumed would be least sensitive to modifications: N- and C- terminal ends of the protein and loop regions, excluding those that are part of CDRs and involved in antigen binding. We analyzed the physicochemical properties and structural orientation of amino acids within the selected regions and proposed ten artificial acceptor sites for N-linked glycosylation in a first glycoengineering campaign (Figure 1A and B). N-X-T glycosylation sequons (with X any amino acid except proline) were introduced at the selected sites by mutation or by the addition of either an N-terminal peptide tag based on a motif frequently occurring in loop D-E of the VHHs, but with an N-glycosylation site introduced (D-D/N-A-N-A-T), or a C-terminal GT1.4tail GlycoTag (Kaup et al., 2011) (sequences in Supplementary A). The variants of GBP designed as such will be referred to as GBP glycovariants.

**Figure 1.**
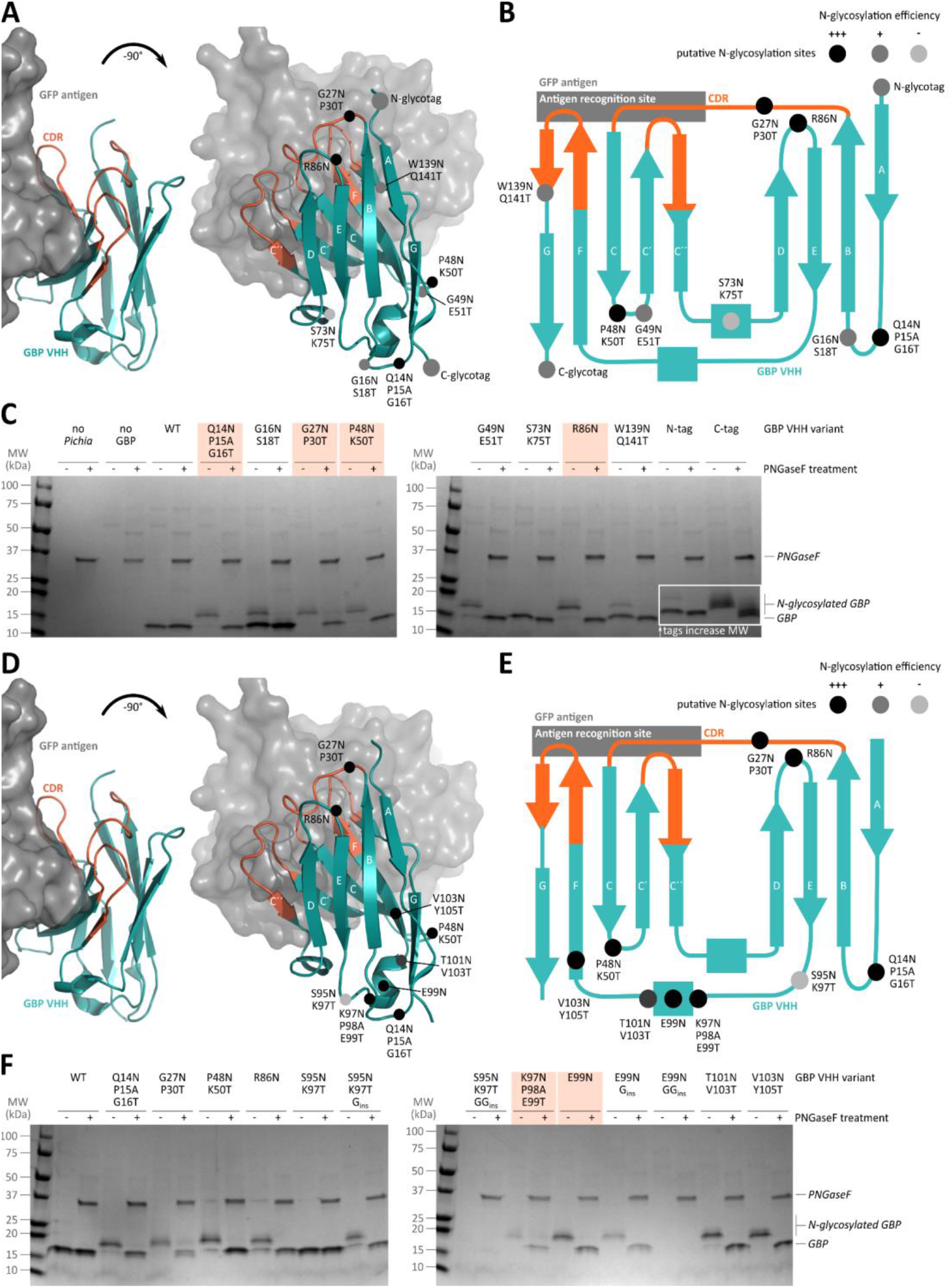
Artificial N-glycosylation introduced in a representative VHH. (A and B) Artificial N-glycosylation at ten sequons introduced in a representative VHH. Eight sites within seven loop regions are indicated in the structure of GFP-binding VHH GBP (PDB id 3OGO (Kubala et al., 2010)) and its topology scheme in which amino acid mutations required for introduction of N-glycosylation sites were introduced. Additionally, N- and C-terminal tags (QADDANATVQLVESGGA and VSSLQAAAAAANATVAAASGDVWDIHHHHHH, respectively) were introduced flanking the VHH coding sequence to introduce an N-glycosylation signature at the N- and C-terminal ends of the protein. (D and E) Additional sites in the E-F loop selected for introduction of N-linked glycosylation signatures are indicated in the structure and topology scheme of GBP. Saturation of the symbol indicates whether the site was more (black), less (dark grey) or not (light grey) efficiently N- glycosylated in VHH GBP as indicated by SDS-PAGE and western blot. Orange indicates CDR-regions of the VHH. (C and F) Coomassie-stained SDS-PAGE of supernatant of *P. pastoris* GlycoSwitchM5 expressing GBP glycovariants (single representative clone per glycovariant shown). Mutations performed to yield a specific variant are indicated; Gins and GGins indicate insertion of one or two glycine residues, respectively. A downwards shift of bands on SDS-PAGE upon PNGaseF treatment indicates removal of an N-glycan. VHH variants selected for further analysis are indicated by orange shading.

Effective therapeutic protein glycotargeting of the macrophage mannose receptor (MMR) requires the presence of exposed mannose residues, preferably in the context of an N-glycan structure that also occurs in humans. Therefore, we produced our VHH glycovariants in the *P. pastoris* GlycoSwitchM5 strain (Jacobs et al., 2009). Glycoproteins produced by this strain are mostly modified with the Man_5_GlcNAc_2_ N-glycan as the most abundant glycoform rather than typical yeast hypermannosylation, also resulting in easily detected discrete bands in SDS-PAGE and more easily deconvoluted ESI mass spectra. Man5GlcNAc2 is one of the highest-affinity natural N-glycans for the MMR (Consortium for Functional Glycomics, 2006).

Supernatants of multiple clones of the GBP glycovariant-expressing GlycoSwitchM5 cultures were either mock treated or treated with the deglycosylating enzyme PNGaseF and subjected to SDS-PAGE analysis. A shift in the molecular weight of the PNGaseF-treated protein confirmed the presence of N-linked glycans on all of the GBP glycovariants except for S73N-K75T (Figure 1C and Supplementary B, representative clone of each glycovariant shown). The glycosylation efficiency (site occupancy) varied between the different VHH variants, suggesting that the N-X-T sequon localization governs glycosylation efficiency.

In a second glycoengineering campaign, we designed a set of GBP glycovariants with N-glycosylation sequons at sites 95, 97, 99, 101 and 103, in the previously untouched E-F loop region (Figure 1D and E). Moreover, as it has been reported that proline (P) residues immediately upstream or downstream of the N-X-T sequon can negatively impact glycosylation efficiency (Bañó-Polo et al., 2011), we generated some variants with ‘extended’ glycosylation sequons (G-N-X-T, GG-N-X-T, N-X-T-G and N-X-T-GG). In these variants, glycine (G) residues were introduced immediately upstream or downstream of the N-X-T sequon to avoid vicinal prolines.

Despite the presence of a structured region of α-helical character within the E-F loop, each of the modified positions could be successfully N-glycosylated (Figure 1F and Supplementary B). Interestingly, the insertion of a single glycine downstream of the S95N-K97T sequon was required to aid N-glycosylation – presumably due to the proline residue at position 98 – whereas the addition of a glycine upstream of E99N was not required for N-glycosylation and even slightly lowered expression levels. Both downstream of S95N-K97T and upstream of E99N, the insertion of two glycines abolished recombinant protein expression.

### The exact position and/or local environment of the N-glycosylation sequon affects site occupancy

The differences in site occupancy (Figure 1C), for example between the Q14N-P15A-G16T and much less efficiently glycosylated G16N-S18T variants – both within loop A-B – and between the P48N-K50T and G49N-E51T variants – both within loop C-C’ – suggest that the exact localization of an N-X-T sequon within a loop of the protein structure is important for N-glycosylation efficiency. Hence, in a third glycoengineering campaign, we performed N-X-T scanning of three selected loops in our benchmark VHH GBP: loop B-C (AHo numbering 27-40), which coincides with CDR1 and contains GBP glycovariant G27N-P30T, loop C-C’ (AHo numbering 47-53), which contains GBP glycovariant P48N-K50T, and loop D-E (AHo numbering 83-88), which contains GBP glycovariant R86N. A first N-X-T scanning set of GBP glycovariants was generated by introducing an N-X-T sequon at every single position within the amino acid stretches 27-40 (loop B-C), 48-52 (within loop C-C’) and 83-88 (loop D-E) by mutation of model VHH GBP, while keeping the loop length constant (Supplementary C-A). A second N-X-T scanning set of glycovariants was generated by peptide inserts at several positions within the selected loops. These were short peptides (N-A-T) consisting of an N-glycosylation sequon only, longer peptides (D-N-A-N-A-T) based on a motif frequently occurring in VHH loop D-E, or single or double glycine residues (G-G) flanking the N-glycosylation sequon if the amino acid preceding it was a proline (Supplementary C-A). Upon production in *P. pastoris* GlycoSwitchM5, VHH glycosylation was evaluated via endoglycosidase treatment and subsequent SDS-PAGE analysis followed either by Coomassie staining or a western blot detecting the C-terminal His6-tag of the VHH (Supplementary C-B and -C). Except for mutant A85N-N87T, which could not be expressed in repeated experiments, all glycovariants were prone to N-glycosylation. Site occupancy varied substantially with the exact position of the sequon (observed in duplicate or triplicate clones per glycovariant). Although the majority of the sites within the selected loop regions showed a high N-glycan site-occupancy (non-glycosylated band barely detectable), some sites (e.g. positions 29 and 46, and a short insert at position 85-86) showed less efficient glycosylation as evidenced by the presence of a lower molecular weight band on SDS-PAGE. Hence, the exact position and/or the local environment of the N glycosylation sequon again affects N-glycosylation site occupancy.

### Characterization of a selected set of N-glycosylated GBP glycovariants

#### Glycan analysis

While it was clear that we had many good sites for further work, we selected the sites in each loop where we had first identified that the native sequence could be mutated to an N-glycosylation sequon, and for which very high site occupancy was observed. Selected glycovariants Q14N-P15A-G16T, G27N-P30T, P48N-K50T, R86N, K97N-P98A-E99T and E99N (shaded orange in Figure 1C and F) were purified for further characterization (Figure 2A). Endoglycosidase treatment and subsequent SDS-PAGE analysis of the purified proteins confirmed the presence of N-linked glycans on the purified glycovariants (Figure 2B and C), but it was also clear that our nickel-NTA and gel filtration purification strategy had biased to the non-glycosylated minor species, in some cases more than others, likely due to protein peaks running across multiple selected fractions, with peak tails not retained for pooling to the final preparation. Hence, the percentage site-occupancy as determined by analysis of purified protein is an underestimation.

**Figure 2.**
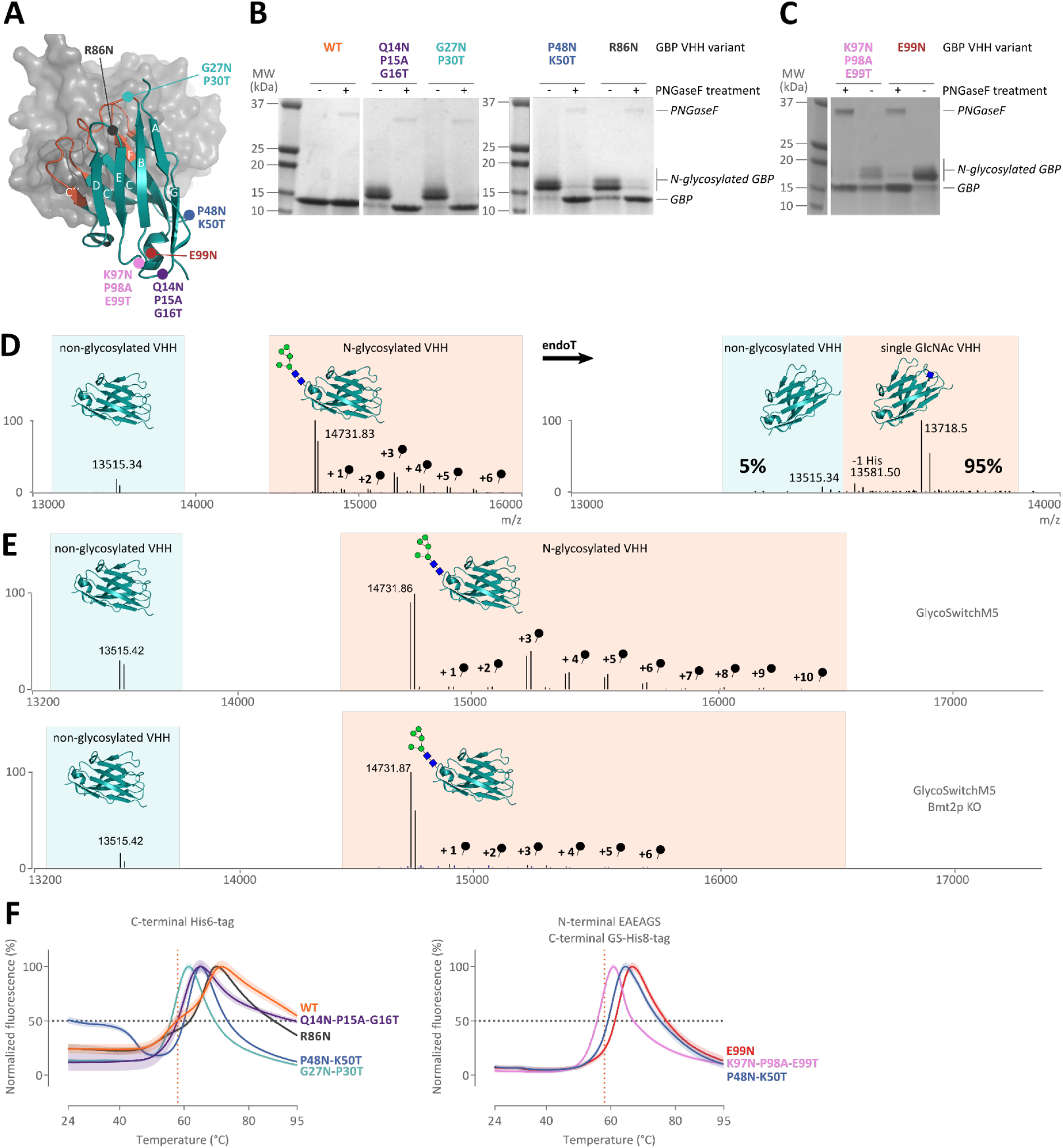
Selected GBP glycovariants are N-glycosylated with high site-occupancy and limited effect on thermal stability. (A) Six GBP glycovariants were selected for *P. pastoris* GlycoSwitchM5 production and IMAC-SEC purification alongside regular GBP (WT). (B) Coomassie-stained SDS-PAGE of GBP and His-6 tagged glycovariants, treated with (+) and without (-) deglycosylating enzyme PNGaseF after denaturation. (C) Coomassie-stained SDS-PAGE of GBP glycovariants that carry an N-terminal EAEAGS sequence (partially processed) and C-terminal GS-His8 tag, treated with (+) and without (-) deglycosylating enzyme PNGaseF after denaturation. (D) Intact protein mass spectrometry of representative VHH GBP-Q14N-P14A-G16T shows a large fraction of N-glycosylated protein and confirms Man_5_GlcNAc_2_ nature of the major N-glycan. Unidentified hexoses are shown as black filled circles. Endoglycosylase T (endoT) digestion of the high-mannose N-glycans to a single GlcNAc allowed for evaluation of the percentage of site-occupancy. (E) Knockout of the β-mannosyltransferase 2 gene (Bmt2p) in *P. pastoris* GlycoSwitchM5 led to a more homogeneous N-glycosylation profile of GBP-Q14N-P15A-G16T. (F) Melting curves of GBP glycovariants. Data points represent mean values of triplicate experiments, standard deviations are indicated by shading.

Intact protein mass spectrometry of purified wildtype GBP (WT) returned a single set of two peaks, indicating the presence of GBP molecules with either an N-terminal glutamine or an N-terminal pyroglutamate (17 Da difference) (Supplementary D). In contrast, the spectra of GBP glycovariants showed an additional cluster of peaks at higher molecular weight. The 1216 Da shift of the major peak corresponds with the presence of a Man_5_GlcNAc_2_ glycan. Small peaks at incremental 162 Da intervals indicate the presence of additional hexoses on the N-glycan (representative example GBP-Q14N-P14A-G16T in Figure 2D; other glycovariants in Supplementary D). Endoglycosidase treatment leaves a single GlcNAc, allowing mass spectrometrical quantification of site occupancy. Peak quantification of endoglycosidase T (endoT) digested purified protein samples demonstrated high N-glycosylation site occupancies from 71% up to 96% (Figure 2D, Supplementary D and E and Table 1), but this variance is largely due to bias towards the non-glycosylated species during purification, as set out above, as the non-glycosylated species in the selected glycovariants were not or barely detectable on SDS-PAGE prior to purification.

**Table 1.** N-glycosylation of selected GBP glycovariants slightly affects biophysical parameters of nanobodies but does not hamper GFP binding. Site occupancy of purified proteins was determined by intact protein mass spectrometry after endoT digestion of the high-mannose N-glycans to a single GlcNAc. Tm was determined as the temperature corresponding to 50% of fluorescence in DSF melting curves. Kinetics parameters K_d_, k_on_ and k_off_ were determined by BLI with immobilized GFP. Mean and standard deviation (SD) of triplicate measures are shown.

| GBP variant | Modifications | | Site occupancy of purified protein (%) | $T_m$ (°C) | | $K_D$ (M) | | $k_{on}$ (M<sup>-1</sup>s<sup>-1</sup>) | | $k_{off}$ (s<sup>-1</sup>) | |
| --- | --- | --- | --- | --- | --- | --- | --- | --- | --- | --- | --- |
|  | N-terminal | C-terminal |  | mean | SD | mean | SD | mean | SD | mean | SD |
| WT | | His6 | N/A | 58.0 | 3.7 | $2.8 \cdot 10^{-10}$ | $3.2 \cdot 10^{-10}$ | $8.1 \cdot 10^5$ | $3.3 \cdot 10^5$ | $1.6 \cdot 10^{-4}$ | $1.2 \cdot 10^{-4}$ |
| Q14N-P15A-G16T | | His6 | 95 | 57.7 | 1.9 | $2.7 \cdot 10^{-10}$ | $3.0 \cdot 10^{-10}$ | $1.1 \cdot 10^6$ | $6.3 \cdot 10^5$ | $1.8 \cdot 10^{-4}$ | $7.2 \cdot 10^{-5}$ |
| G27N-P30T | | His6 | 96 | 55.8 | 0.2 | $2.8 \cdot 10^{-10}$ | $3.4 \cdot 10^{-10}$ | $1.0 \cdot 10^6$ | $5.0 \cdot 10^5$ | $1.7 \cdot 10^{-4}$ | $1.2 \cdot 10^{-4}$ |
| P48N-K50T | | His6 | 96 | 59.9 | 0.3 | $1.5 \cdot 10^{-10}$ | $6.2 \cdot 10^{-11}$ | $1.3 \cdot 10^6$ | $7.3 \cdot 10^5$ | $1.7 \cdot 10^{-4}$ | $2.6 \cdot 10^{-5}$ |
| R86N | | His6 | 82 | 61.7 | 0.6 | $5.1 \cdot 10^{-11}$ | $4.1 \cdot 10^{-11}$ | $1.7 \cdot 10^6$ | $2.9 \cdot 10^5$ | $7.9 \cdot 10^{-5}$ | $5.4 \cdot 10^{-5}$ |
| P48N-K50T | EAEAGS | GS-His8 | 94 | 59.1 | 0.4 | $7.8 \cdot 10^{-11}$ | $3.4 \cdot 10^{-11}$ | $6.0 \cdot 10^5$ | $3.9 \cdot 10^5$ | $4.2 \cdot 10^{-5}$ | $3.4 \cdot 10^{-5}$ |
| K97N-P98A-E99T | EAEAGS | GS-His8 | 74 | 55.7 | 0.2 | $7.2 \cdot 10^{-11}$ | $4.9 \cdot 10^{-11}$ | $9.9 \cdot 10^5$ | $1.0 \cdot 10^6$ | $4.0 \cdot 10^{-5}$ | $1.6 \cdot 10^{-5}$ |
| E99N | EAEAGS | GS-His8 | 71 | 61.3 | 0.4 | $1.1 \cdot 10^{-10}$ | $1.4 \cdot 10^{-10}$ | $8.6 \cdot 10^5$ | $2.3 \cdot 10^5$ | $9.4 \cdot 10^{-5}$ | $1.0 \cdot 10^{-4}$ |

N-glycan profiling of glycovariants Q14N-P15A-G16T, G27N-P30T and P48N-K50T through capillary electrophoresis on a DNA sequencer confirmed that in addition to the main Man_5_GlcNAc_2_, higher molecular weight glycans were present in small quantities (Supplementary F and G). These additional N-glycans (with a main peak of Hex_8_GlcNAc_2_) were recalcitrant to α-1,2-mannosidase and were only partially hydrolyzed by Jack Bean α-1,2/3/6-mannosidase. This behavior is identical to that of glycans observed earlier on other *P. pastoris* GlycoSwitchM5-produced proteins, where we identified the Hex_8_GlcNAc_2_ species as Man_5_GlcNAc_2_ with a substituent consisting of Manβ-1,2-Manβ-1,3-Glcα-1,3-R (Laukens et al., 2020). It is known that the β-mannosyltransferase gene family is responsible for the addition of β-mannose residues in *P. pastoris* (Mille et al., 2008). Blocking the synthesis of these apparently also allows for the underlying α-Glc residue to be hydrolyzed by the secretory system α-glucosidase. Indeed, disruption of the *BMT2* gene by inducing a double-guide CRISPR-Cpf1 mediated deletion resulted in a more homogeneous N-glycosylation profile of GBP-Q14N-P14A-G16T (Figure 2E), consistent with a previous report that Bmt2p is the key β-mannosyltransferase in the synthesis of these off-target N-glycans (Hopkins et al., 2011).

#### Molecular dynamics

To evaluate whether the presence of glycans would not impede VHH-antigen interactions, we initially used the GlycoSHIELD approach (Gecht et al., 2021) to model VHH surface accessibility. Upon grafting conformation libraries of either the main GlycoSwitchM5 glycan Man_5_GlcNAc_2_ (Supplementary H) or the regular yeast Golgi-type Man_9_GlcNAc_2_ N-glycan (Figure 3A) to each of our six preferred N-glycosylation sites in the modified crystallographic structure of GBP (Kubala et al., 2010), the surface closely around those N-glycosylation sites was partially shielded based on calculated solvent accessibility using 1.4 to 10 Å probes (Figure 3B and Supplementary H).

**Figure 3.**
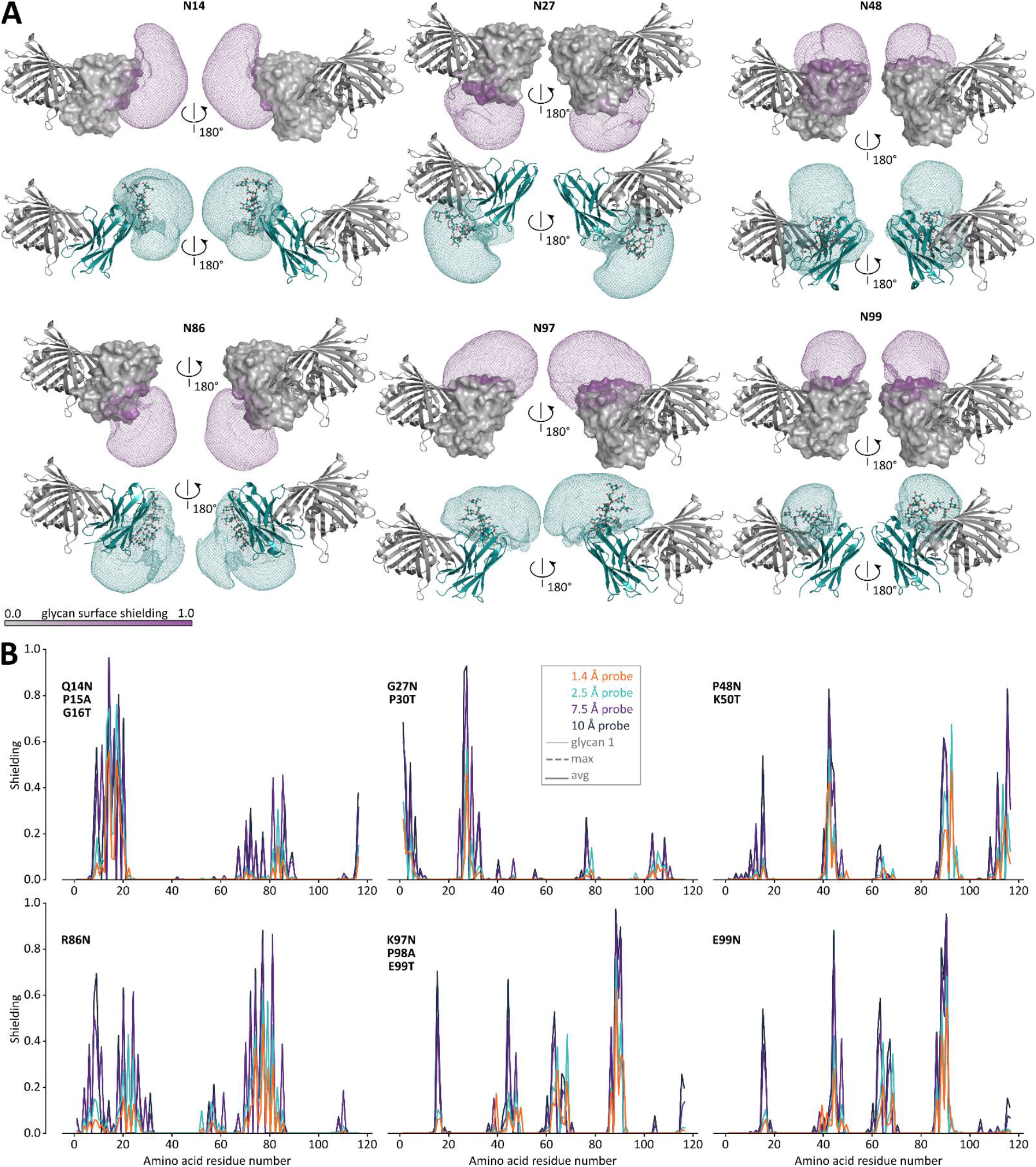
Molecular dynamics and GlycoSHIELD models of each of the selected single N-glycosylation site glycovariants. (A) GlycoSHIELD Man9GlcNAc2 conformation libraries (purple mesh) grafted into glycosylation sites Q14N, G27N, P48N, R86N, K97N and E99N in a modified structure of GBP with its antigen GFP (grey, PDB id 3OGO; Kubala et al., 2010). Shielding of the protein surface (shown in grey-to-purple surface gradient) was determined after calculating the solvent accessible surface area using the GlycoSHIELD script (Gecht et al., 2021) with a 10.0 Å probe. Molecular dynamics simulations of the selected GBP single N-glycosylation site glycovariants in explicit solvent are represented below as cyan ‘cartoon’ ribbons, with glycan structures represented in ball-and-stick mode. The 0.5% space occupancy isocontours of a yeast Golgi-type Man_9_GlcNAc_2_ for N-glycosylation sites N14, N27, N48, N86, N97 and N99 are shown as cyan mesh. (B) GlycoSHIELD Man_9_GlcNAc_2_ shielding of the protein surface using four different probe sizes of 1.4 (orange), 2.5 (cyan), 7.5 (purple), and 10.0 (dark blue) Å.

**Figure 4.**
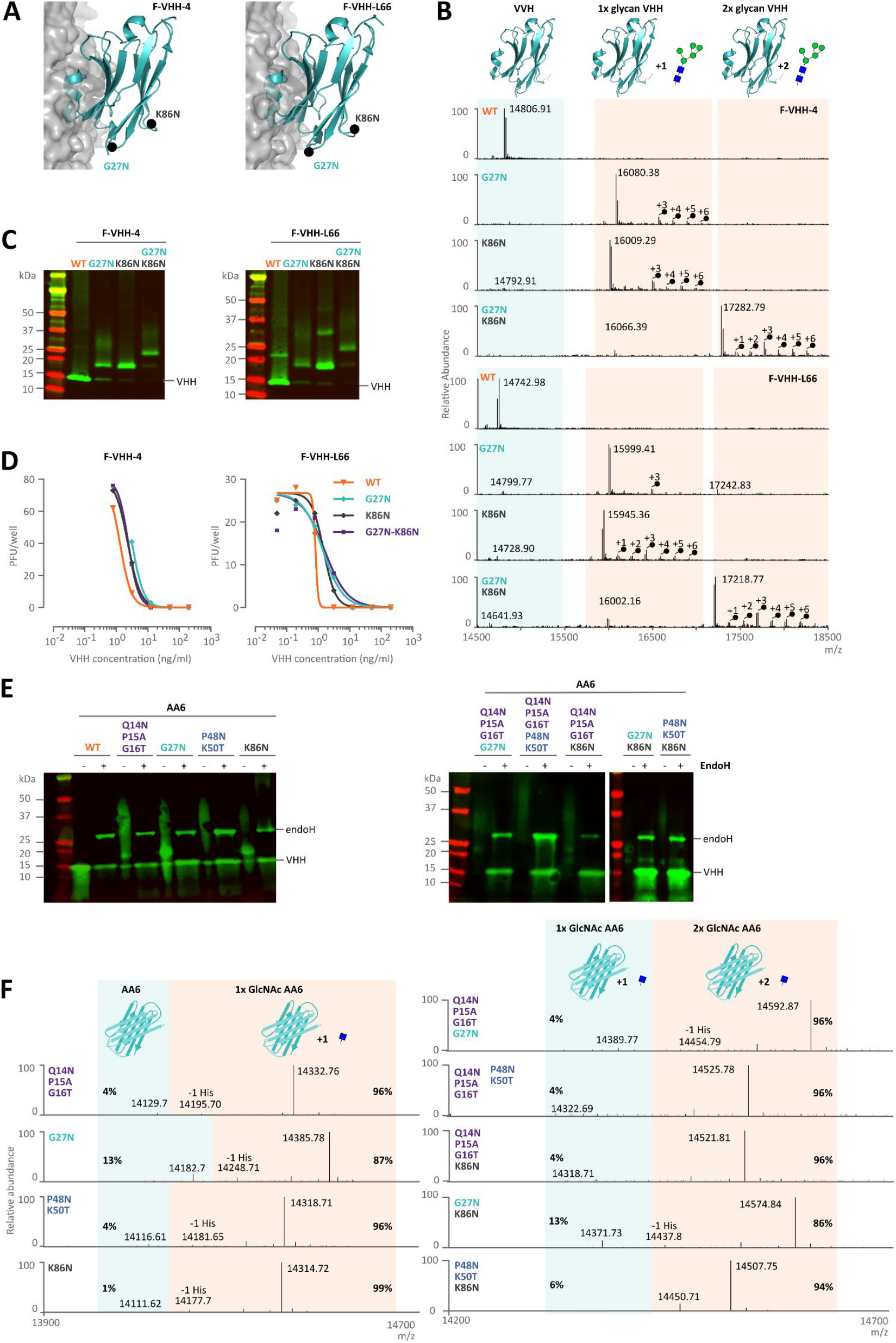
The N-glycosylation approach for GBP could be extrapolated to other VHHs. (A) N-glycosylation sites introduced in F-VHH- 4 and F-VHH-L66 are indicated in their structures (PDB id 5TOJ and 5TOK (Rossey et al., 2017)). VHHs in cyan, RSV fusion protein in prefusion conformation in grey. (B) Intact protein mass spectrometry and (C) DyLight800 anti-His tag western blot detection of supernatants of *P. pastoris* GlycoSwitchM5 expressing F-VHH-4 and F-VHH-L66 and the G27N, K86N and G27N-K86N-glycovariants thereof. Black filled circles indicate additional (162 Da) hexoses. (D) Neutralization of respiratory syncytial virus by F-VHH-4 and F-VHH-L66 glycovariants. (E) Supernatants of *P. pastoris* expressing AA6 wildtype (WT) and glycovariants G14N-P15A-G16T, G27N, P48N-K50T and K86N, and glycovariants carrying two N-glycosylation sequons in a single construct. A downwards shift of bands on SDS-PAGE upon Endoglycosidase H (endoH) treatment indicates removal of N-glycans. Western blot with DyLight800 anti-His tag detection. (F) Intact mass spectrometry peak height analysis after EndoH endoglycosidase digest allows quantification of glycovariant site occupancy.

To gain more detailed insight in the conformational space occupied by the selected N-glycans while considering the possibility of protein-glycan interactions, we resorted to 2 μs molecular dynamics simulations of the glycoprotein in explicit solvent. Our preferred six selected N-X-T sites were each introduced in the structure of GBP and derivatized *in silico* with a Man_9_GlcNAc_2_ N-glycan. Subsequent molecular dynamics showed the sections of space around the VHH that the glycans can sample, indicating that such glycan substitutions could be used to shield off access to most areas of the VHH surface except for – intentionally – the CDRs, which remain unoccluded even when modeling these large Man_9_GlcNAc_2_ N-glycans (Figure 3A and Supplementary I).

#### Affinity

To evaluate whether VHH antigen binding is indeed retained upon N-glycosylation, we assessed affinity of wildtype GBP and the six glycovariants, using biolayer interferometry. Wildtype GBP displays nanomolar affinity for its antigen GFP (Kubala et al., 2010). In line with the molecular modeling results, the presence of an N-glycan on the glycovariants did not significantly alter antigen binding kinetics (Table 1 and Supplementary J).

#### Stability

Glycovariant stability was assessed via differential scanning fluorimetry (Lo et al., 2004) (Figure 2F). The melting curve of wildtype GBP showed a two-step unfolding process starting at 45 °C. For the glycovariants, we observed both one- and two-step unfolding kinetics with the melting onset around the same temperature. Importantly, typical VHH stability was retained in all glycovariants, with T_m_ values varying from 55.7 to 61.6 °C. Of note is that in repeated measurements, His6-tagged GBP glycovariant P48N-K50T displayed high fluorescence at low temperature (Figure 2F, left). Binding of the fluorescent dye to the low-temperature conformation suggested exposure of (some) hydrophobic residues, possibly due to mutation of a structurally important proline residue in the C-C’ loop. However, upon expression of the same glycovariant in a format that differs by carrying an N-terminal incompletely processed EAEAGS leader sequence and a C-terminal His8 rather than His6 tag, native state dye binding decreased (Figure 2F, right). This suggests that the aberrant behavior of this mutant is not an inevitable consequence of the glycosylation mutations. That being said, this loop contains key residues that have evolved to keep the VHH fold stable and highly soluble in the absence of a light chain (Nguyen et al., 2000), and its mutation is often avoided also in VHH sequence humanization for this reason.

#### Glyco-engineering is transferrable to other VHHs

To evaluate whether our findings can be extrapolated to other (therapeutically relevant) VHHs, we introduced a selection of N-linked glycosylation sequons in the RSV prefusion F (fusion) protein-specific VHHs F-VHH-4 and F-VHH-L66 (Rossey et al., 2017) and in AA6, a VHH binding *Clostridium difficile* exotoxin (Feng et al., 2017; Yang et al., 2014).

Upon introduction of N-glycosylation sequons at positions 27 and/or 86 in the RSV-specific VHHs (Figure 3A), intact protein mass spectrometry and anti-His western blot of the *P. pastoris* GlycoSwitchMan5 supernatant revealed that the glycoengineered F-VHH-4 and F-VHH-L66 VHHs were modified with one (in case of one N-glycosylation sequon) or two (in case of two N-glycosylation sequons) oligomannose-type N-glycans, with near-complete site-occupancy (Figure 3B and C). Moreover, the presence of (predominantly) Man_5_GlcNAc_2_ N-glycans did not preclude RSV neutralization (Figure 3D).

Wildtype *P. pastoris* production of glycoengineered AA6 variants showed that single- or double-site N-glycosylation was successful in this VHH as well (Figure 3E), though very heterogeneous glycosylation was observed, typical of yeast hypermannosylation. Only at sites 27 and 86, a significant portion of the glycans was of the yeast ‘core type’. These results highlight once again that GlycoSwitch engineering of *P. pastoris* is critical if more homogeneous N-glycosylation is desired. Upon endoglycosidase-removal of the highly heterogeneous N-glycans leaving a single GlcNAc, site occupancies of 87% to 99% were observed at sites 14, 27, 48 and 86 of AA6 and similarly high double-site occupancies were seen upon combining two of these N-glycosylation sequons (Figure 3F).

#### GlycoSwitchMan5-produced VHHs for macrophage phagosome delivery

Next, we wanted to evaluate if the addition of GlycoSwitchMan5-produced N-glycans could potentiate VHH uptake by macrophages. To this end, GBP-WT and the G27N-P30T and R86N glycovariants were conjugated with a fluorescent AlexaFluor488 (AF488) probe via NHS ester chemistry. Using spectrophotometry, the degree of labeling was determined to be 0.6, 1.0 and 1.2 mol/mol for the WT, G27N-P30T and R86N GBP variants, respectively. This was confirmed by capturing the labeled VHHs in a dilution series on a nickel-coated plate and quantifying fluorescence (Supplementary K).

To test macrophage uptake of the GBP variants, co-cultures of macrophages (Mf4/4 cells) and control cells (HEK293T cells) were incubated with PBS, AF488-labeled GBP-WT, or AF488-labeled GBP-G27N-P30T for 40 minutes, and subsequently analyzed by flow cytometry. Analysis revealed that all Mf4/4 cells (CD45^+^), but not the HEK293T cells (CD45^-^), were able to capture or take up GBP-G27N-P30T in a concentration dependent manner (Figure 5A, lower panels). In contrast, neither Mf4/4 nor HEK293T cells were able to bind GBP-WT (Figure 5A, upper panels). These observations demonstrate that the engineered GlycoSwitchMan5 N-glycans on GBP allow efficient binding and/or uptake of the VHH by macrophages. Confocal microscopy was used to determine whether the glycovariants are actually taken up by the macrophages, or if they merely bind to the macrophage cell surface. Figure 5B illustrates that both the G27N-P30T and R86N GBP glycovariants are readily internalized by macrophages, with a small part of the internalized GBP glycovariants overlapping with LysoTracker-stained lysosomes. In contrast, the vast majority of macrophages did not take up GBP-WT. To confirm uptake of the GBP glycovariants by macrophages and to understand the kinetics of this process, live cell imaging of Mf4/4 cells was performed upon addition of GBP via perfusion. This analysis revealed that uptake of the GBP glycovariants, but not of GBP-WT, could be detected already a few minutes after administration (stills in Figure 5C and videos in Supplementary L).

**Figure 5.**
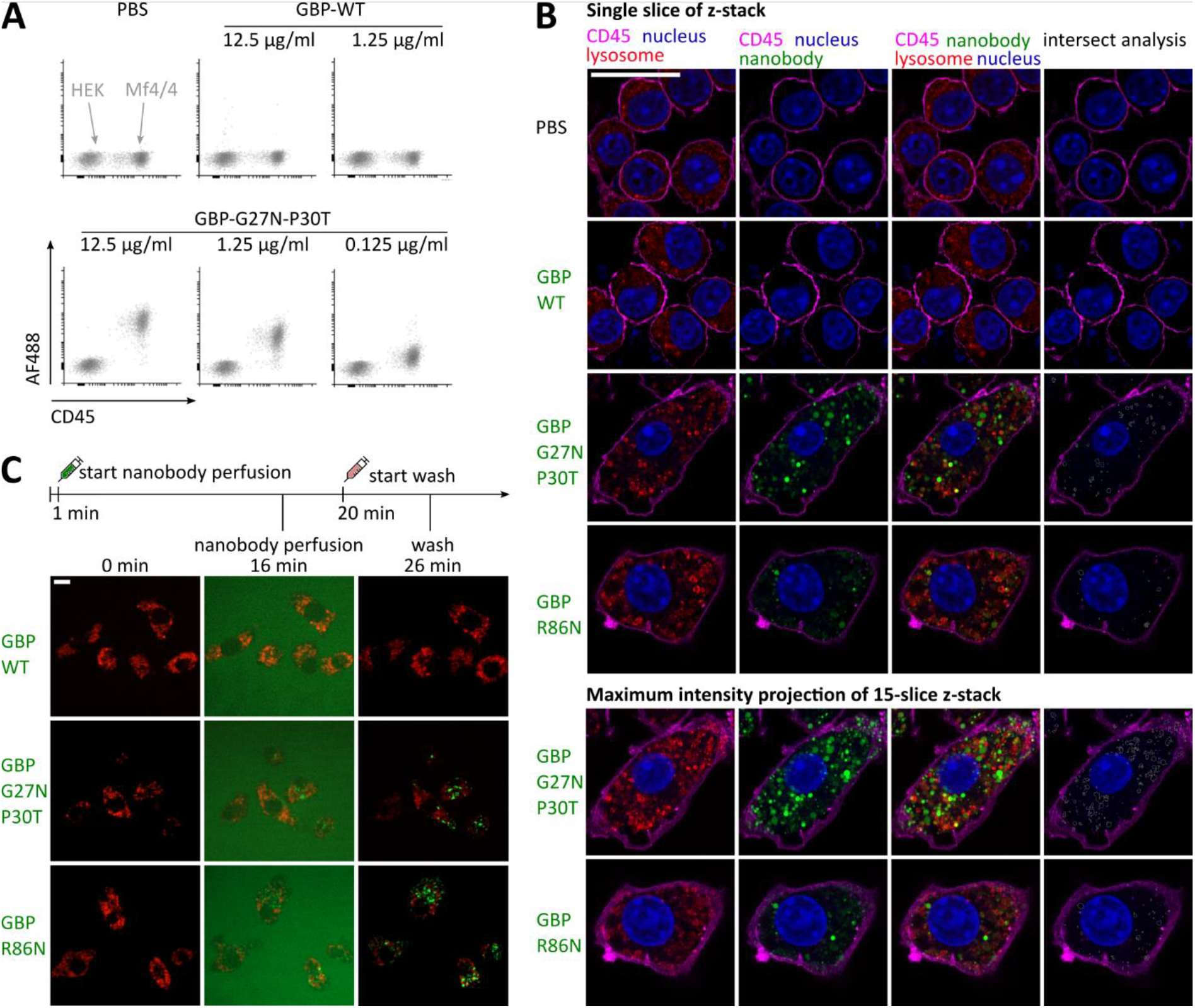
Glycan-mediated VHH internalization by Mf4/4 macrophages. (A) A co-culture of HEK293T and Mf4/4 cells was inoculated with AlexaFluor488 (AF488)-labeled benchmark VHH GBP or GlycoSwitchMan5-expressed GBP-G27N-P30T. Uptake was measured via flow cytometry. (B) Confocal images of immunofluorescence stained Mf4/4 cells. Cells were incubated with PBS, or AF488-labeled GBP-WT or GlycoSwitchMan5 glycovariants GBP-G27N-P30T and GBP-R86N. Lysosomes were visualized by addition of LysoTracker (red). AF647-coupled antibody against CD45 was used to stain macrophage membranes (magenta). Cell nuclei were identified by DAPI staining (blue). An intersect analysis (grey) identified intersection between the lysosomal and VHH populations. White scaling bar indicates 10 μm. (C) Stills from live cell imaging of GBP glycovariant uptake by Mf4/4 cells. Mf4/4 macrophages stained with LysoTracker (red) were incubated at 37 °C in 5% CO_2_. 12.5 μg of AF488-labeled GBP or GBP glycovariant (green) was perfused at 1 ml/min. After 20 minutes of incubation, VHHs were washed out by perfusion of culture medium at 0.1 ml/min and images were recorded for 10 more minutes. White scaling bars indicate 10 μm.

Finally, we set out to investigate if Man_5_GlcNAc_2_-modified GBP is also efficiently taken up by macrophages *in vivo.* Therefore, PBS or 12.5 μg of either GBP-WT or GBP-G27N-P30T was administered to the lungs of BALB/C mice. One hour after administration, bronchoalveolar lavage fluid (BALF) was collected to investigate the uptake of GBP by alveolar macrophages. Flowcytometric analysis revealed that, as expected, none of the CD45^-^F4/80^-^ cells had taken up detectable levels of GBP. In contrast, the majority of CD45^+^F4/80^+^ alveolar macrophages had taken up GBP-WT and GBP-G27N-P30T. In agreement with the *in vitro* uptake assay, GBP- G27N-P30T was taken up by alveolar macrophages over 10-fold more efficiently than GBP-WT (Figure 6A and B). To confirm that the GBP glycovariants were taken up by the alveolar macrophages, part of the BALF was used for confocal microscopy. After allowing the alveolar macrophages to adhere, they were fixed and stained for CD45 and F4/80, present on alveolar macrophages. Indeed, GlycoSwitchMan5-modified GBP but not GBP-WT (likely below detection limit) could be clearly visualized inside these alveolar macrophages (Figure 6C).

**Figure 6.**
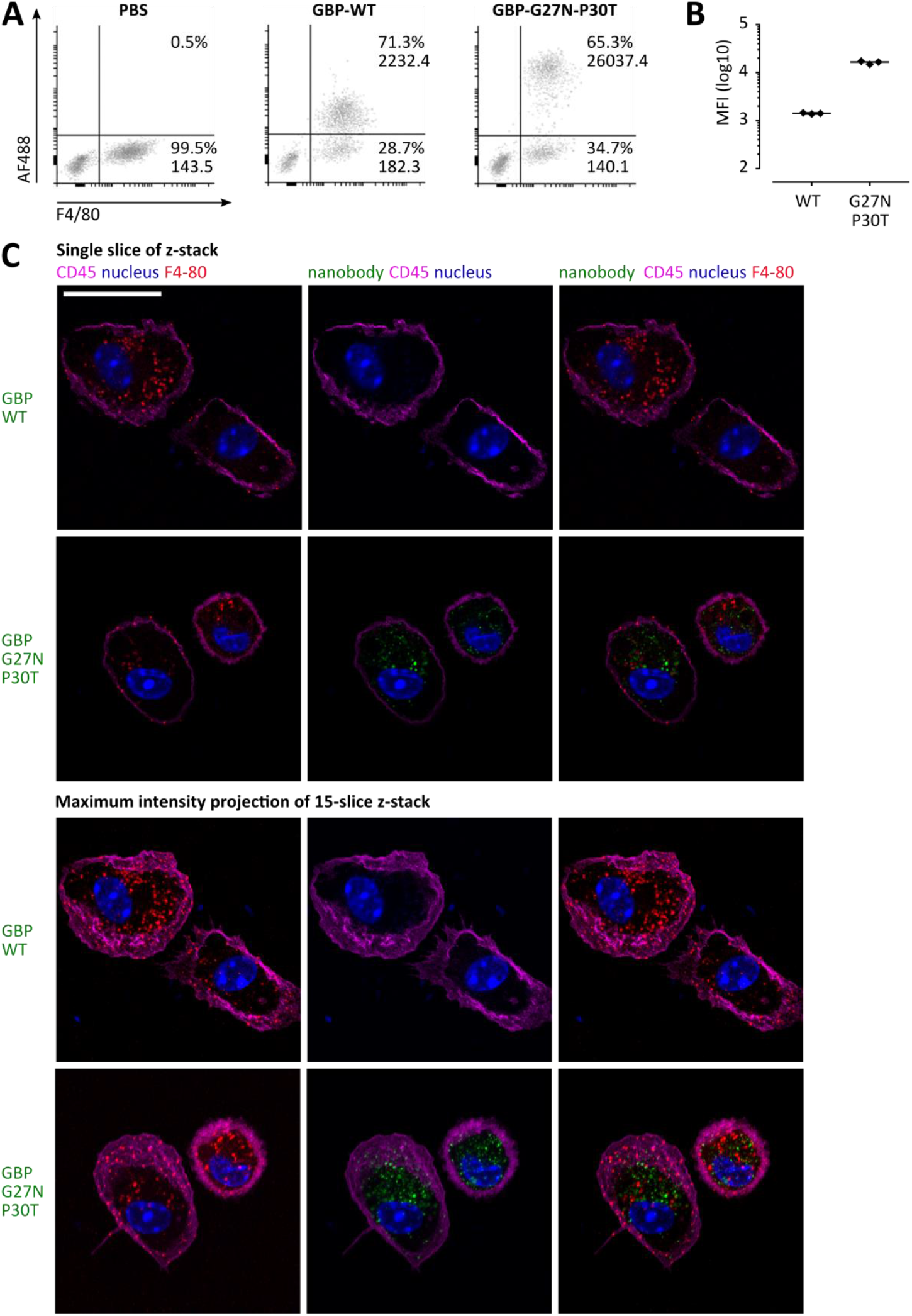
*In vivo* glycan-mediated VHH internalization by alveolar macrophages. (A) Flow cytometric analysis of BALF cells from BALB/C mice that had PBS (left) or 10 μg of AF488-labeled GBP-WT (middle) or GlycoSwitchMan5-expressed GBP-G27N-P30T (right) administered to the lungs. The numbers in the right quadrants represent the % of alveolar macrophages (F4/80+) that had (upper right quadrant) or had not (lower right quadrant) taken up AF-488 labeled GBP-WT or GlycoSwitchMan5 GBP-G27N-P30T and their respective AF488 median fluorescence intensity (MFI). (B) Flow cytometric analysis of BALF cells of BALB/C mice that had 10 μg of AF488-labeled GBP-WT or GlycoSwitchMan5 GBP-G27N-P30T administered to the lungs. The graph shows the AF488 MFI of CD45+ alveolar macrophages from three mice treated with GBP-WT or GlycoSwitchMan5 GBP-G27N-P30T. (C) Confocal microscopy of BALF cells. Alveolar macrophages were fixed and stained with CD45 and F4/80, present on alveolar macrophages. Cell nuclei were identified by DAPI staining (blue). White scaling bar indicates 10 μm. Both a single image and the maximum intensity projection of a 15-slide z-stack are shown.

## Discussion

In this study, we aimed to expand the VHH bioengineering toolbox by enrichment of the available chemical space in the molecule, via the introduction of N-glycans. In a structure-guided approach, VHHs were engineered to allow N-linked glycosylation at sites where the presence of an N-glycan was not expected to interfere with VHH folding or target recognition; in addition, three loop regions were screened exhaustively via N-X-T scanning, and some sites were flanked with glycine residues to avoid the influence of vicinal prolines. Upon expression in *P. pastoris,* most VHH variants showed at least partial N-linked glycosylation, with a few exceptions. No N-glycosylation could be detected on the S73N-K75T and S95N-K97T variants of our benchmark VHH GBP; limited accessibility of the site for the oligosaccharyltransferase or the importance of the local secondary structure in folding may explain this observation. We also found that the A85N-N87T, S95N-K97T-GGins and GGins-E99N variants of GBP were not expressed at all, which suggests that the specific mutations introduced here may have disturbed the local structure, leading to folding or stability issues. Amongst over fifteen glycovariants with very high site occupancy (non-glycosylated band undetected on SDS-PAGE), three were also described in the patent literature as this project was already ongoing (Q14N-P15A-G16T, P48N-K50T, and V103N-Y105T; Buyse & Boutton, 2016) whereas the others are first reported here. Further studies showed that a GlycoSwitchMan5 N-glycan at positions 14, 27, 48, 86, 97 and 99 did not compromise VHH stability or target binding, with possible exception of a sequence context-dependent destabilization of the P48N-K50T mutant that hits the VHH-typical FR2 framework region that was evolved to accommodate the lack of a light chain in these antibodies. Molecular dynamics simulations suggest that glycan chains at all of these positions are projected away from the antigen-binding region, minimizing the risk of interfering with antigen recognition, which was experimentally confirmed for several glycovariants of anti-GFP and anti-RSV VHHs. In a final set of experiments, we showed that GlycoSwitchMan5-type N-glycans at the select positions that we tested can potentiate macrophage-specific uptake both *in vitro* and *in vivo.*

Targeted N-linked glycosylation of VHHs now unlocks a range of opportunities for modulation of protein characteristics and glycan-based targeting. For instance, N-linked glycans on the VHH scaffold may shield proteolytically sensitive peptide stretches in protease-rich environments like the gut. Moreover, glycosylation may provide a solution for therapeutic VHHs that show immunogenicity issues by shielding off B-cell epitopes, and it is a proven technological option for increasing hydrodynamic radius of small proteins, thereby strongly enhancing circulatory half-life by reduced renal clearance (Koury, 2003).

Targeted introduction of N-linked glycans also paves the way for site-specific conjugation of the VHH. The introduced glycans can be used as a bio-orthogonal handle for conjugation of half-life extending moieties, drugs and tracers. This provides an alternative or complementary strategy to the classic conjugation approaches on primary amines or cysteines, which suffer from limited site-control and often lead to heterogeneous product profiles. Glycan-based conjugation strategies have been successfully used in the production of antibody-drug conjugates (ADCs), and resulted in a more homogeneous product profile (Van Delft et al., 2014; Van Delft et al., 2015a; Van Delft et al., 2015b). We are reporting on this application of glyco-VHHs elsewhere (Van Breedam et al., 2021).

Finally, N-linked glycans can also be utilized for cell- and/or tissue-specific targeting of the VHHs. In the human body, glycans can interact with carbohydrate-binding proteins called lectins. Many of these lectins are membrane-associated proteins, and they are typically expressed in a cell- and/or tissue-specific manner, which makes them attractive targets in cell-/tissue-specific glycan-based targeting. The glycan specificity of lectins is determined by the nature and the valency of their carbohydrate recognition domains, and most lectins show a marked preference for specific mono/oligosaccharides in a particular presentation. Consequently, specific glycan types give access to (a set of) specific cell types, and production of a glyco-engineered VHH in expression systems with different N-glycosylation phenotypes, gives access to different cell populations. As we have shown, production of a glyco-engineered VHH in a yeast strain that produces Man_5_GlcNAc_2_-enriched N-glycans allows targeting of the VHH for endocytosis by the macrophage mannose receptor (MMR), a macrophage-restricted lectin that traffics to the phagosomal compartment. In this way, a phagolysosomal targeting chimeric molecule is also easily obtained. We are looking forward to explore the utility of this newly enabled VHH engineering in the treatment of various disease indications. For instance, pathogens like *Listeria, Brucella, Mycobacterium tuberculosis* and *Acetinobacter baumanni* are able to infect macrophages and survive and propagate within the phagosomal compartment. Diseases caused by such pathogens are typically hard to combat with biopharmaceuticals, as these have to be delivered to the macrophage phagolysosome. It will be interesting to investigate whether high-mannose glycan-based targeting of antimicrobial VHHs could significantly lower the effective therapeutic dose or increase the therapeutic index, similar to what was achieved with an antibody-antibiotic conjugate against *S. aureus* (Lehar et al., 2015).

The current study provides a blueprint for the targeted introduction of N-linked glycans in the VHH scaffold yielding glycoVHHs. Targeted glycosylation of VHH provides a new and extra level of versatility for this important drug class.

## Supporting information

GlycoVHH_Supplementary

## Grant numbers

- ERC Consolidator Grant: grant no. 616966, “GlycoTarget”,
- UGent Geconcerteerde onderzoeksacties (GOA): no grant number
- FWO Strategisch Basis Onderzoek (SBO): grant no. S007817N, “GlycoDelete”

- Funded W.V.B., F.S., K.T. and S.V.
- FWO Regulier Onderzoeksproject: grant no. G041417N

- Funded A.V.H.
- IWT Strategisch Basisonderzoek (SB): grant no. 131545

- Funded L.v.S.

## Acknowledgements

We thank Eef Parthoens from the VIB BioImaging Core for technical (live cell) imaging assistance.

## Supplemental material is supplied in a separate file

