## Supplementary material for "GlycoVHH: Introducing N-glycans on the camelid VHH antibody scaffold - Optimal sites and use for macrophage delivery": GlycoVHH_Supplementary

### Supplemental material

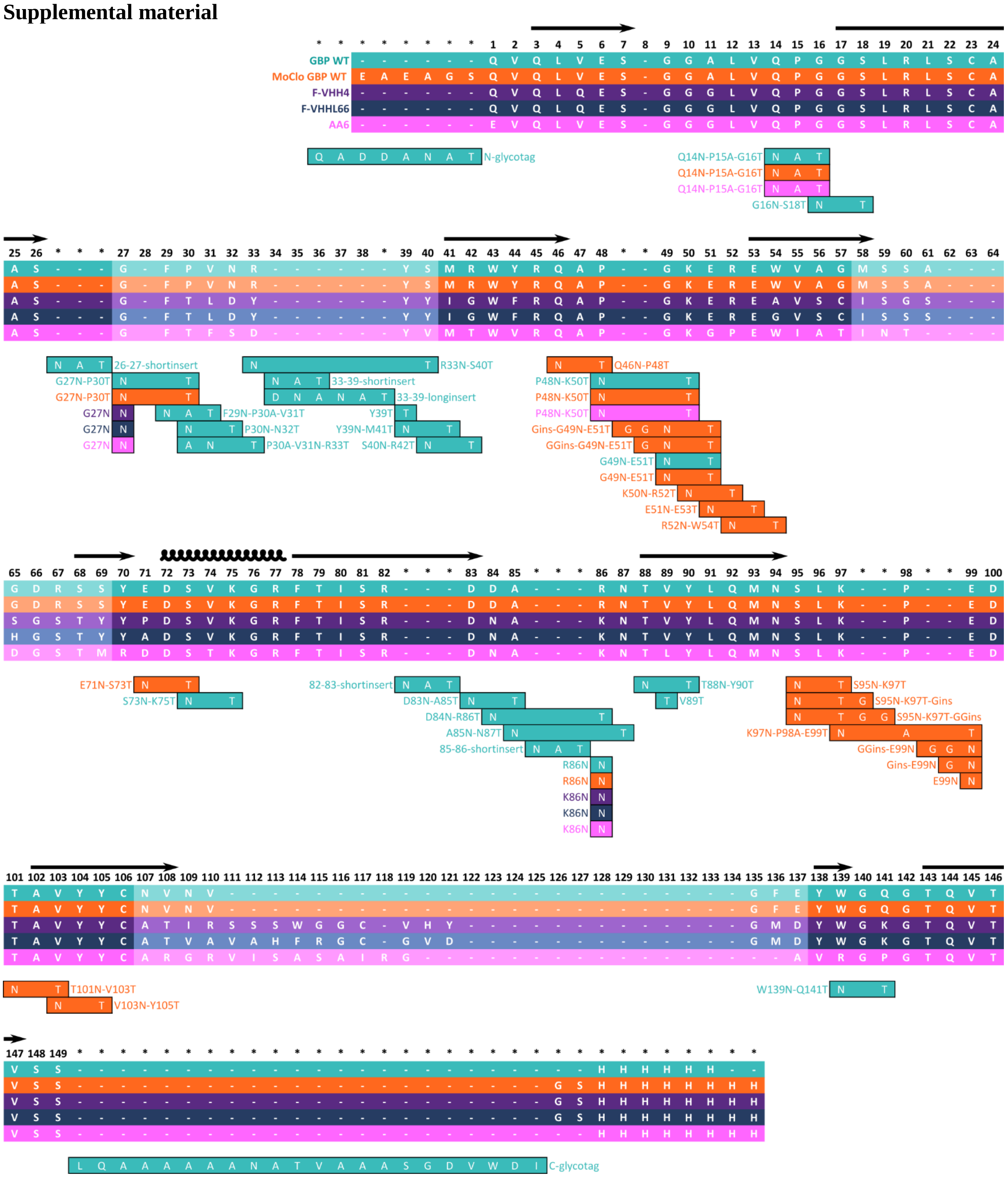


**Supplementary A. Aligned amino acid sequences of VHHs and their glycovariants.** Amino acid sequences of GBP (regular: cyan, MoClo: orange), F-VHH4 (purple), F-VHHL66 (blue) and AA6 (pink) are numbered according to AHo numbering conversion (Honegger and Plückthun, 2001) and CDR regions are shaded in a lighter hue. The amino acid mutations required to insert N-glycosylation sequons in each VHH are depicted below the sequences as blocks in corresponding colors, and tagged by the name of the glycovariant containing the mutations. Secondary structure elements of GBP are indicated as described by Kubala et al (2010) (arrows: β-sheets; curls: α-helices).

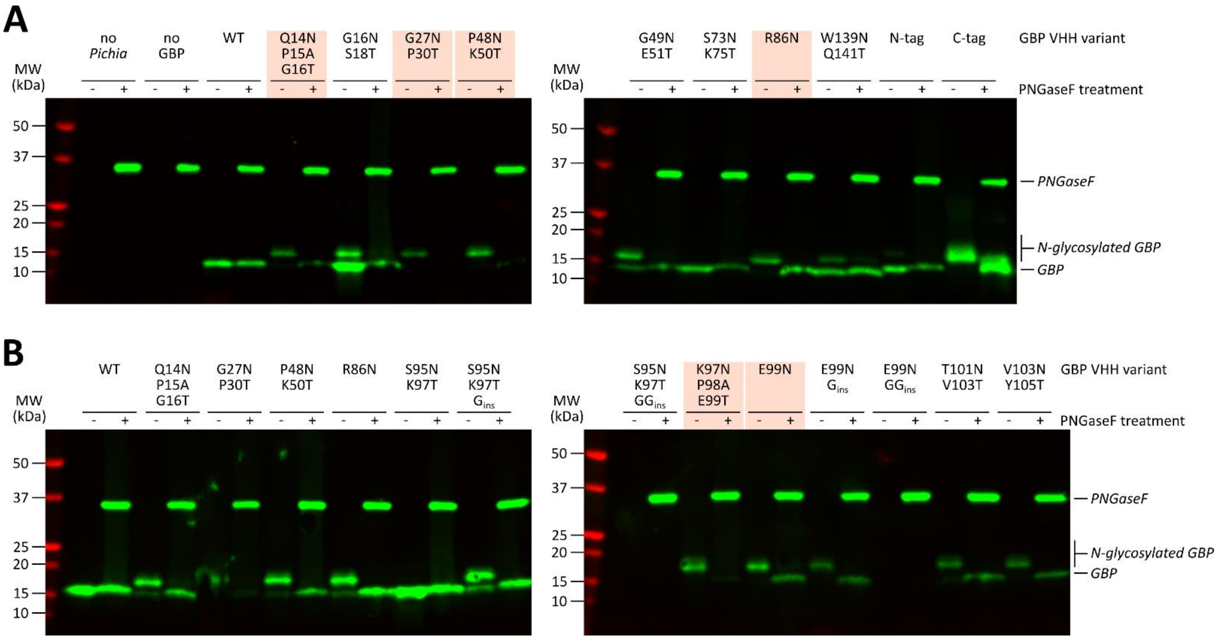


**Supplementary B. Artificial N-glycosylation introduced in a representative VHH (western blot validation).** Artificial N-glycosylation at (A) ten sequons introduced in a representative VHH, and (B) additional sites in the E-F loop. DyLight800-tagged anti-His western blot of supernatant of *P. pastoris* GlycoSwitchM5 expressing GBP glycovariants (single representative clone per glycovariant shown). Mutations performed to yield a specific variant are indicated; Gins and GGins indicate insertion of one or two glycine residues, respectively. A downwards shift of bands on SDS-PAGE upon PNGaseF treatment indicates removal of an N-glycan. VHH variants selected for further analysis are indicated by orange shading.

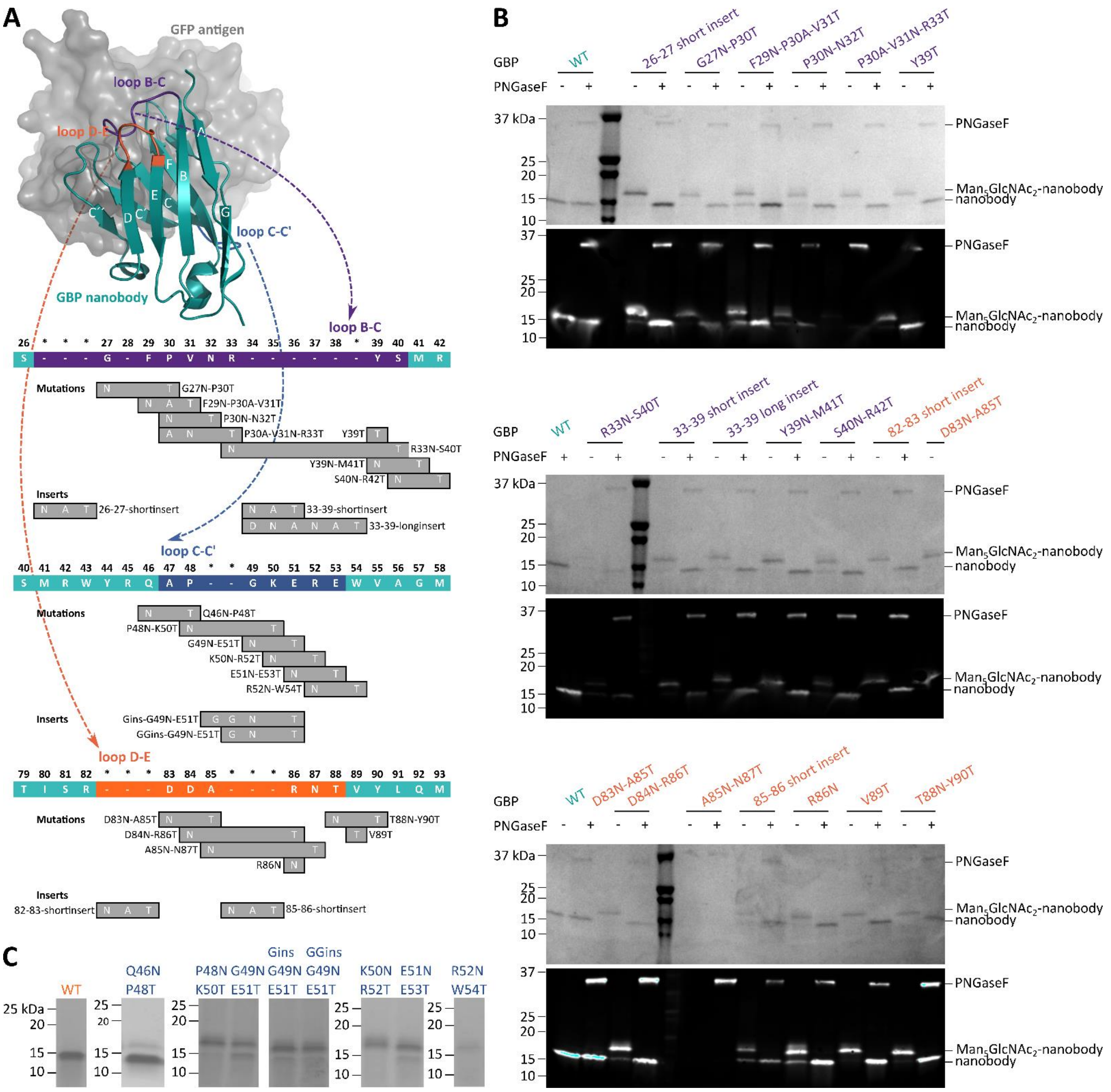


**Supplementary C. N-X-T scanning of selected loops of the benchmark GBP single-domain antibody (VHH).** (A) Loops B-C, C-C' and D-E of GFP-binding VHH GBP were selected for N-X-T scanning. Artificial N-glycosylation sites were introduced by site-specific mutation or insertion as indicated in the scheme. (B) N-glycosylation of loops B-C and D-E was successful with varying site occupancy at all sites except position 85. Coomassie-stained SDS-PAGE and anti-His western blot of *P. pastoris* GlycoSwitchM5 supernatant expressing GBP-WT (wildtype, unmodified) and glycovariants, treated with (+) and without (-) deglycosylating enzyme PNGaseF after denaturation. (C) N-glycosylation of loop C-C' was successful with highly varying site occupancy. Coomassie-stained SDS-PAGE of *P. pastoris* GlycoSwitchM5 supernatant expressing GBP-WT (wildtype, unmodified) and glycovariants.

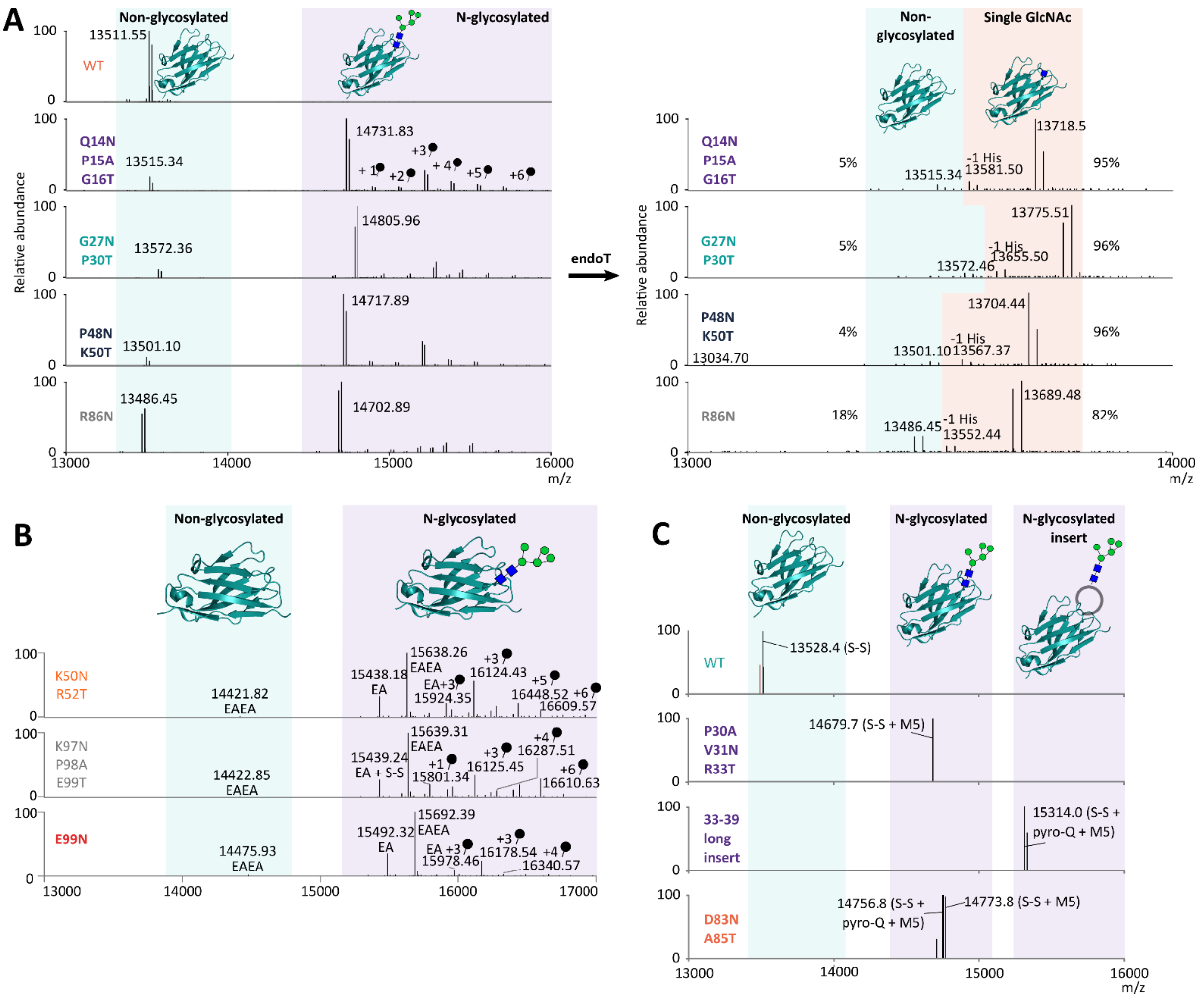


**Supplementary D. GBP glycovariants are N-glycosylated with varying site-occupancy (additional data not shown in main figures).** (A) Intact protein mass spectrometry of selected purified GBP glycovariants shows varying fractions of N-glycosylated protein and confirms $Man_5GlcNAc_2$ nature of the major N-glycan. Endoglycosylase T (endoT) digestion of the high-mannose N-glycans to a single GlcNAc allowed for evaluation of the percentage of site-occupancy. (B) Intact protein mass spectrometry of *P. pastoris* GlycoSwitchM5 supernatant expressing GBP glycovariants that carry an N-terminal EAEAGS sequence (partially processed) and C-terminal GS-His8 tag. (B) Intact protein mass spectrometry of *P. pastoris* GlycoSwitchM5 supernatant expressing GBP glycovariants in B-C and D-E loops. Peaks representing non-glycosylated VHHs are shaded in cyan, peaks representing N-glycosylated VHHs in purple, peaks representing endoT-cleaved N-glycosylated VHHs in orange. Additional hexoses are indicated by black circles. EAEA or EA: indicates presence of N-terminal Glu-Ala-repeats, pyro-Q: N-terminal pyroglutamine instead of glutamine.

| | Variant | Monoisotopic Mass | Sum Intensity | Full sum intensity | Fraction (%) |
|---|---|---|---|---|---|
| GBP-WT | no peaks except for the naked protein and its pyroglutamine form; 0% N-glycosylation | | | | |
| GBP-Q14N-P15A-G16T | 0x GlcNAc/pyroglutamine/-His | 13378 | 1.18E+04 | | |
| | 0x GlcNAc/-His | 13395 | 3.24E+03 | | |
| | 0x GlcNAc/pyroglutamine/ | 13515 | 1.46E+05 | | |
| | 0x GlcNAc | 13532 | 5.86E+04 | 2.20E+05 | 5.4 |
| | 1x GlcNAc/pyroglutamine/-His | 13581 | 2.50E+05 | | |
| | 1x GlcNAc/-His | 13598 | 1.30E+05 | | |
| | 1x GlcNAc/pyroglutamine/ | 13718 | 2.26E+06 | | |
| | 1x GlcNAc | 13735 | 1.21E+06 | 3.85E+06 | 94.6 |
| GBP-G27N-P30T | 0x GlcNAc/pyroglutamine/ | 13572.48 | 1.84E+05 | | |
| | 0x GlcNAc | 13589.48 | 1.23E+05 | 3.07E+05 | 4.0 |
| | 1x GlcNAc/pyroglutamine/-2His | 13501.48 | 2.26E+04 | | |
| | 1x GlcNAc/-2His | 13518.48 | 3.81E+04 | | |
| | 1x GlcNAc/pyroglutamine/-His | 13638.48 | 2.78E+05 | | |
| | 1x GlcNAc/-His | 13655.48 | 3.66E+05 | | |
| | 1x GlcNAc/pyroglutamine/ | 13775.48 | 2.88E+06 | | |
| | 1x GlcNAc | 13792.48 | 3.79E+06 | 7.37E+06 | 96.0 |
| GBP-P48N-K50T | 0x GlcNAc/pyroglutamine/ | 13501.4 | 8.37E+04 | | |
| | 0x GlcNAc | 13518.4 | 3.39E+04 | 1.18E+05 | 3.6 |
| | 1x GlcNAc/pyroglutamine/-His | 13567 | 1.30E+05 | | |
| | 1x GlcNAc/-His | 13584 | 6.08E+04 | | |
| | 1x GlcNAc/pyroglutamine/ | 13704 | 1.99E+06 | | |
| | 1x GlcNAc | 13721 | 9.73E+05 | 3.15E+06 | 96.4 |
| GBP-R86N | 0x GlcNAc/pyroglutamine/-His | 13332 | 5.74E+04 | | |
| | 0x GlcNAc/-His | 13349 | 6.25E+04 | | |
| | 0x GlcNAc/pyroglutamine/ | 13469 | 8.16E+05 | | |
| | 0x GlcNAc | 13486 | 8.54E+05 | 1.79E+06 | 18.2 |
| | 1x GlcNAc/pyroglutamine/-His | 13535 | 2.94E+05 | | |
| | 1x GlcNAc/-His | 13552 | 3.08E+05 | | |
| | 1x GlcNAc/pyroglutamine/ | 13672 | 3.50E+06 | | |
| | 1x GlcNAc | 13689 | 3.94E+06 | 8.04E+06 | 81.8 |
| GBP-P48N-K50T | 0x GlcNAc | 14481 | 7.00E+00 | 7.00E+00 | 5.4 |
| EAEAGS/GS-His8 | 1x GlcNAc/-His | 14547 | 2.30E+01 | | |
| | 1x GlcNAc | 14684 | 1.00E+02 | 1.23E+02 | 94.6 |
| GBP-K97N-P98A-E99T | 0x GlcNAc/-His | 14286 | 9.00E+00 | | |
| EAEAGS/GS-His8 | 0x GlcNAc | 14423 | 3.50E+01 | 4.40E+01 | 26.5 |
| | 1x GlcNAc/-His | 14489 | 2.20E+01 | | |
| | 1x GlcNAc | 14626 | 1.00E+02 | 1.22E+02 | 73.5 |
| GBP-E99N | 0x GlcNAc/-His | 14339 | 1.40E+01 | | |
| EAEAGS/GS-His8 | 0x GlcNAc | 14476 | 4.00E+01 | 5.40E+01 | 28.9 |
| | 1x GlcNAc/-His | 14542 | 3.30E+01 | | |
| | 1x GlcNAc | 14679 | 1.00E+02 | 1.33E+02 | 71.1 |

**Supplementary E. N-glycosylation quantification of selected purified GBP and AA6 glycovariants.** Intact protein mass spectrometry peak quantification. Proteins were treated with endoT or endoH enzyme overnight to reduce N-glycans to a single GlcNAc residue. "Pyroglutamine" indicates pyroglutamine formation of the N-terminal glutamate residue; "x GlcNAc" indicates the number of single GlcNAc residues remaining after EndoH or EndoH cleavage of the high mannose glycan; "-His" indicates the loss of a C-terminal histidine residue.

| | Variant | Monoisotopic Mass | Sum Intensity | Full sum intensity | Fraction (%) |
|---|---|---|---|---|---|
| **AA6-Wt** | 0x GlcNAc/pyroglutamine/-His | 13972 | 1.92E+04 | | |
| | 0x GlcNAc/-His | 13989 | 3.67E+05 | | |
| | 0x GlcNAc/pyroglutamine/ | 14109 | 2.64E+05 | | |
| | 0x GlcNAc | 14126 | 4.28E+06 | 4.93E+06 | 100.0 |
| **AA6-14N** | 0x GlcNAc | 14129 | 2.63E+05 | 2.63E+05 | 3.9 |
| | 1x GlcNAc/-His | 14195 | 1.08E+05 | | |
| | 1x GlcNAc/pyroglutamine/ | 14315 | 2.35E+05 | | |
| | 1x GlcNAc | 14332 | 6.15E+06 | 6.49E+06 | 96.1 |
| **AA6-27N** | 0x GlcNAc/-His | 14045 | 1.05E+05 | | |
| | 0x GlcNAc/pyroglutamine/ | 14165 | 4.34E+04 | | |
| | 0x GlcNAc | 14182 | 1.29E+06 | 1.44E+06 | 13.1 |
| | 1x GlcNAc/pyroglutamine/-His | 14231 | 2.96E+04 | | |
| | 1x GlcNAc/-His | 14248 | 6.94E+05 | | |
| | 1x GlcNAc/pyroglutamine/ | 14368 | 4.01E+05 | | |
| | 1x GlcNAc | 14385 | 8.47E+06 | 9.60E+06 | 86.9 |
| **AA6-48N** | 0x GlcNAc/pyroglutamine/ | 14099 | 3.07E+04 | | |
| | 0x GlcNAc | 14116 | 3.31E+05 | 3.61E+05 | 4.4 |
| | 1x GlcNAc/-His | 14181 | 1.73E+05 | | |
| | 1x GlcNAc/pyroglutamine/ | 14301 | 3.57E+05 | | |
| | 1x GlcNAc | 14318 | 7.41E+06 | 7.94E+06 | 95.7 |
| **AA6-86N** | 0x GlcNAc/-His | 13974 | 0.00E+00 | 1.23E+05 | 1.3 |
| | 0x GlcNAc/pyroglutamine/ | 14094 | 0.00E+00 | | |
| | 0x GlcNAc | 14111 | 1.23E+05 | | |
| | 1x GlcNAc/pyroglutamine/-His | 14160 | 9.10E+03 | | |
| | 1x GlcNAc/-His | 14177 | 4.47E+05 | | |
| | 1x GlcNAc/pyroglutamine/ | 14297 | 3.95E+05 | | |
| | 1x GlcNAc | 14314 | 8.37E+06 | 9.22E+06 | 98.7 |

**Supplementary E (continued). N-glycosylation quantification of selected purified GBP and AA6 glycovariants.** Intact protein mass spectrometry peak quantification. Proteins were treated with endoT or endoH enzyme overnight to reduce N-glycans to a single GlcNAc residue. "Pyroglutamine" indicates pyroglutamine formation of the N-terminal glutamate residue; "x GlcNAc" indicates the number of single GlcNAc residues remaining after EndoH or EndoH cleavage of the high mannose glycan; "-His" indicates the loss of a C-terminal histidine residue.

| | Variant | Monoisotopic Mass | Sum Intensity | Full sum intensity | Fraction (%) |
|---|---|---|---|---|---|
| AA6-14N/27N | 0x GlcNAc/-His | 14048 | 0.00E+00 | | |
| | 0x GlcNAc/pyroglutamine/ | 14168 | 0.00E+00 | | |
| | 0x GlcNAc | 14185 | 2.61E+04 | 2.61E+04 | 0.4 |
| | 1x GlcNAc/pyroglutamine/-His | 14235 | 0.00E+00 | | |
| | 1x GlcNAc/-His | 14252 | 0.00E+00 | | |
| | 1x GlcNAc/pyroglutamine/ | 14372 | 0.00E+00 | | |
| | 1x GlcNAc | 14389 | 2.61E+05 | 2.61E+05 | 3.8 |
| | 2x GlcNAc/pyroglutamine/-His | 14438 | 0.00E+00 | | |
| | 2x GlcNAc/-His | 14455 | 6.33E+04 | | |
| | 2x GlcNAc/pyroglutamine/ | 14575 | 2.13E+05 | | |
| | 2x GlcNAc | 14592 | 6.31E+06 | 6.59E+06 | 95.8 |
| AA6-14N/48N | 0x GlcNAc/-His | 13982 | 0.00E+00 | | |
| | 0x GlcNAc/pyroglutamine/ | 14102 | 0.00E+00 | | |
| | 0x GlcNAc | 14119 | 0.00E+00 | 0.00E+00 | 0.0 |
| | 1x GlcNAc/pyroglutamine/-His | 14168 | 0.00E+00 | | |
| | 1x GlcNAc/-His | 14185 | 0.00E+00 | | |
| | 1x GlcNAc/pyroglutamine/ | 14305 | 1.73E+04 | | |
| | 1x GlcNAc | 14322 | 3.67E+05 | 3.85E+05 | 4.3 |
| | 2x GlcNAc/pyroglutamine/-His | 14371 | 0.00E+00 | | |
| | 2x GlcNAc/-His | 14388 | 4.93E+04 | | |
| | 2x GlcNAc/pyroglutamine/ | 14508 | 2.91E+05 | | |
| | 2x GlcNAc | 14525 | 8.28E+06 | 8.62E+06 | 95.7 |
| AA6-14N/86N | 0x GlcNAc/-His | 13978 | 0.00E+00 | | |
| | 0x GlcNAc/pyroglutamine/ | 14098 | 0.00E+00 | | |
| | 0x GlcNAc | 14115 | 0.00E+00 | 0.00E+00 | 0.0 |
| | 1x GlcNAc/pyroglutamine/-His | 14164 | 0.00E+00 | | |
| | 1x GlcNAc/-His | 14181 | 0.00E+00 | | |
| | 1x GlcNAc/pyroglutamine/ | 14301 | 0.00E+00 | | |
| | 1x GlcNAc | 14318 | 4.02E+05 | 4.02E+05 | 3.8 |
| | 2x GlcNAc/pyroglutamine/-His | 14367 | 0.00E+00 | | |
| | 2x GlcNAc/-His | 14384 | 1.44E+05 | | |
| | 2x GlcNAc/pyroglutamine/ | 14504 | 4.89E+05 | | |
| | 2x GlcNAc | 14521 | 9.67E+06 | 1.03E+07 | 96.2 |
| AA6-27N/86N | 0x GlcNAc/-His | 14030 | 0.00E+00 | | |
| | 0x GlcNAc/pyroglutamine/ | 14150 | 0.00E+00 | | |
| | 0x GlcNAc | 14167 | 4.72E+04 | 4.72E+04 | 0.7 |
| | 1x GlcNAc/pyroglutamine/-His | 14217 | 0.00E+00 | | |
| | 1x GlcNAc/-His | 14234 | 3.67E+04 | | |
| | 1x GlcNAc/pyroglutamine/ | 14354 | 2.81E+04 | | |
| | 1x GlcNAc | 14371 | 8.82E+05 | 9.46E+05 | 13.1 |
| | 2x GlcNAc/pyroglutamine/-His | 14420 | 0.00E+00 | | |
| | 2x GlcNAc/-His | 14437 | 2.85E+05 | | |
| | 2x GlcNAc/pyroglutamine/ | 14557 | 2.69E+05 | | |
| | 2x GlcNAc | 14574 | 5.70E+06 | 6.26E+06 | 86.3 |
| AA6-48N/86N | 0x GlcNAc/-His | 13963 | 0.00E+00 | | |
| | 0x GlcNAc/pyroglutamine/ | 14083 | 0.00E+00 | | |
| | 0x GlcNAc | 14100 | 0.00E+00 | 0.00E+00 | 0.0 |
| | 1x GlcNAc/pyroglutamine/-His | 14150 | 0.00E+00 | | |
| | 1x GlcNAc/-His | 14167 | 0.00E+00 | | |
| | 1x GlcNAc/pyroglutamine/ | 14287 | 7.00E+04 | | |
| | 1x GlcNAc | 14304 | 6.78E+05 | 7.48E+05 | 6.0 |
| | 2x GlcNAc/pyroglutamine/-His | 14353 | 0.00E+00 | | |
| | 2x GlcNAc/-His | 14370 | 2.13E+05 | | |
| | 2x GlcNAc/pyroglutamine/ | 14490 | 6.59E+05 | | |
| | 2x GlcNAc | 14507 | 1.08E+07 | 1.16E+07 | 94.0 |

**Supplementary E (continued). N-glycosylation quantification of selected purified GBP and AA6 glycovariants.** Intact protein mass spectrometry peak quantification. Proteins were treated with endoT or endoH enzyme overnight to reduce N-glycans to a single GlcNAc residue. "Pyroglutamine" indicates pyroglutamine formation of the N-terminal glutamate residue; "x GlcNAc" indicates the number of single GlcNAc residues remaining after EndoH or EndoH cleavage of the high mannose glycan; "-His" indicates the loss of a C-terminal histidine residue.

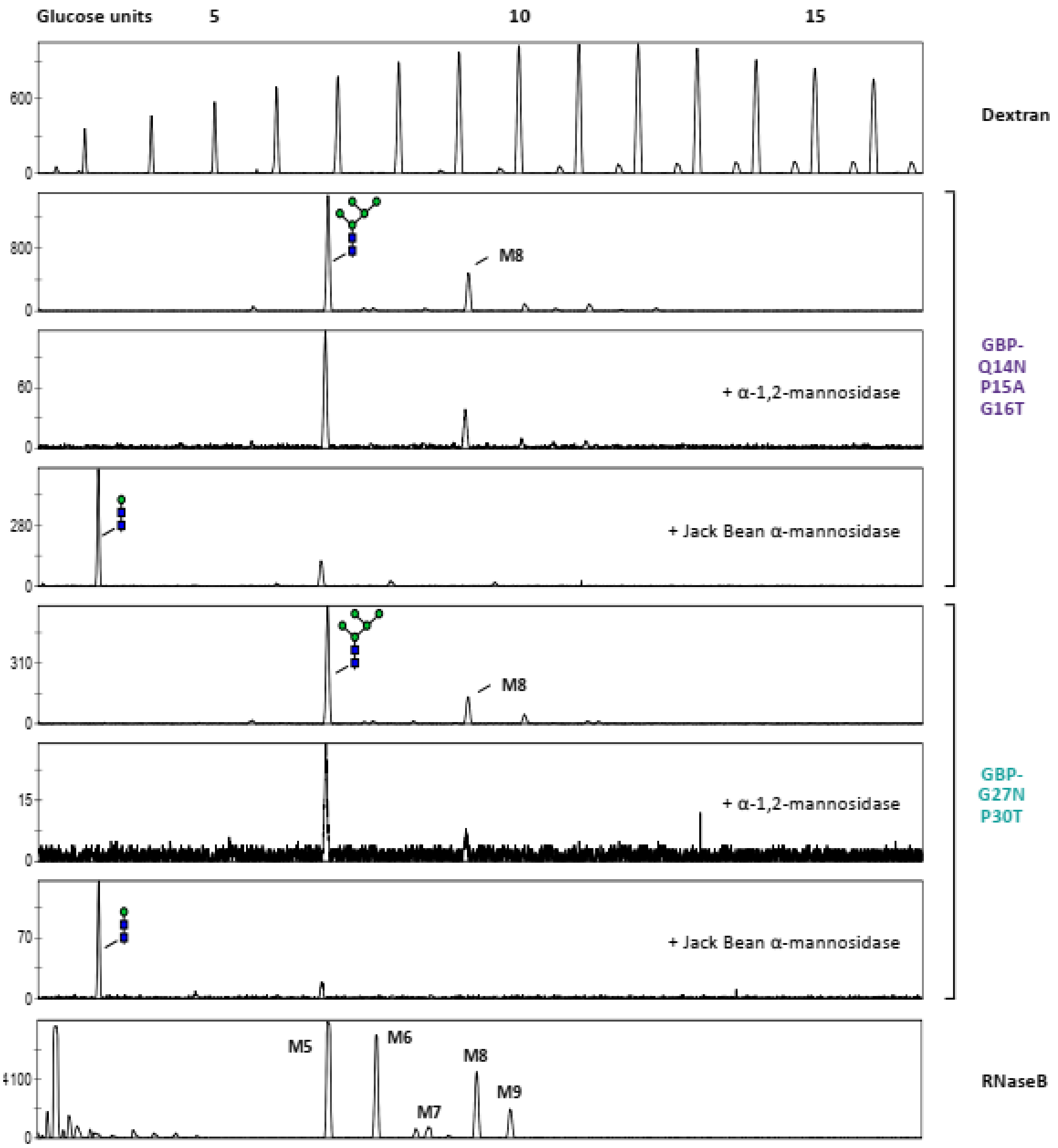


**Supplementary F. DSA-FACE glycan profiling of VHH GBP glycovariants Q14N-P15A-G16T and G27N-P30T.** N-glycans were released from GBP glycovariants by PNGaseF treatment, derivatized with an APTS label (10 mM, in-house produced) and characterized via DNA sequencer-assisted fluorophore-assisted carbohydrate electrophoresis (DSA-FACE) on an ABI3130 Genetic Analyzer (Applied Biosystems). As internal references, partially heat-hydrolyzed *Leuconostoc mesenteroides* dextran and N-glycans from bovine pancreas RNaseB were included. Exoglycosidase digests were performed overnight in 5 mM $NH_4OAc$ pH 5.2 on labeled glycans, with *H. jecorina* α-1,2-mannosidase and Jack Bean α-1,2/3/6-mannosidase.

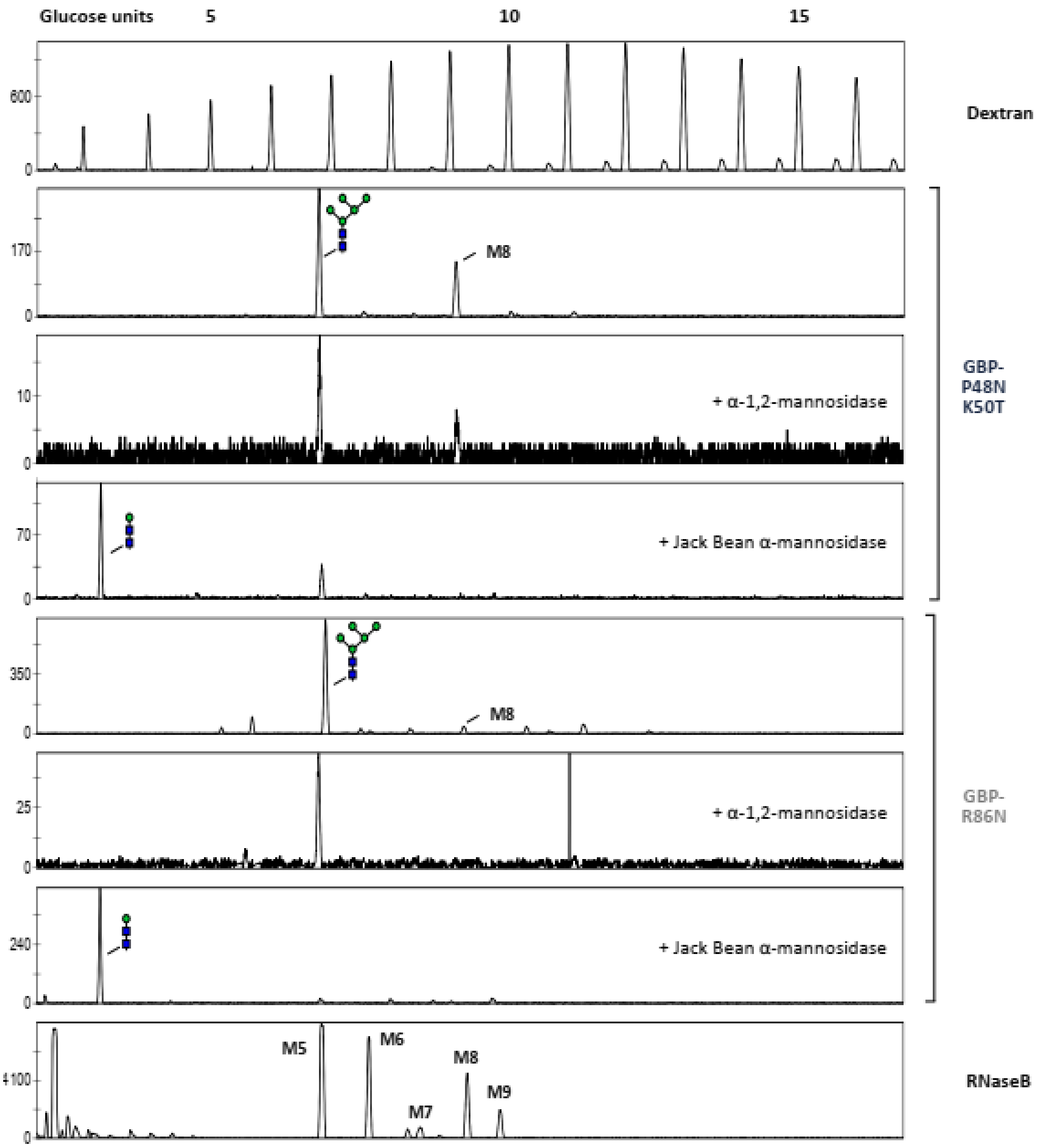


**Supplementary G. DSA-FACE glycan profiling of VHH GBP glycovariants P48N-K50T and R86N.** N-glycans were released from GBP glycovariants by PNGaseF treatment, derivatized with an APTS label (10 mM, in-house produced) and characterized via DNA sequencer-assisted fluorophore-assisted carbohydrate electrophoresis (DSA-FACE) on an ABI3130 Genetic Analyzer (Applied Biosystems). As internal references, partially heat-hydrolyzed *Leuconostoc mesenteroides* dextran and N-glycans from bovine pancreas RNaseB were included. Exoglycosidase digests were performed overnight in 5 mM $NH_4OAc$ pH 5.2 on labeled glycans, with *H. jecorina* α-1,2-mannosidase and Jack Bean α-1,2/3/6-mannosidase.

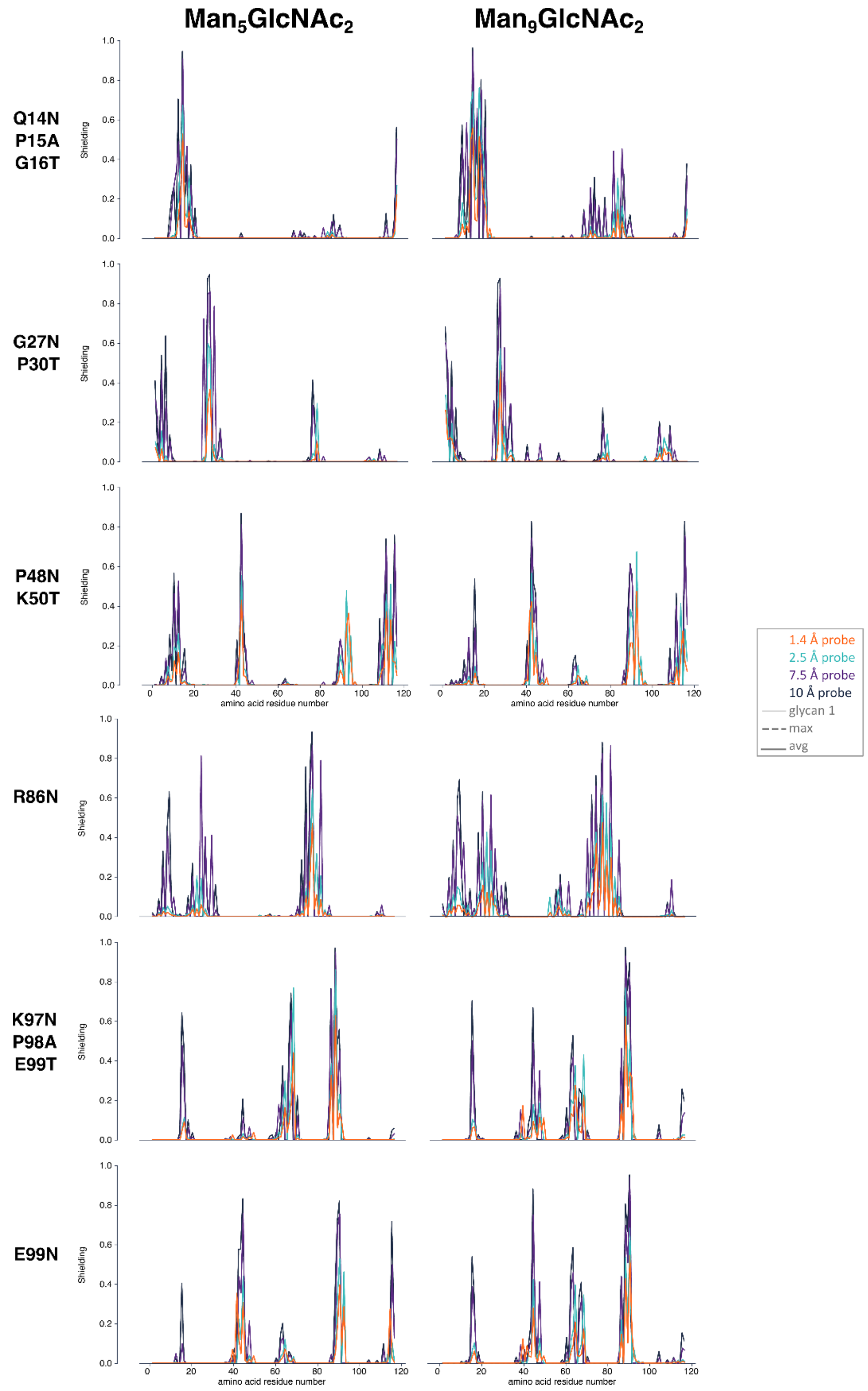


**Supplementary H**. GlycoSHIELD analysis of the selected single N-glycosylation site glycovariants with a Man5GlcNAc2 or Man9GlcNAc2 glycan. Shielding of the protein surface was calculated after calculating the solvent accessible surface area using the GlycoSHIELD script (Gecht et al., 2021) with four different probe sizes of 1.4 (orange), 2.5 (cyan), 7.5 (purple), and 10.0 (dark blue) Å.

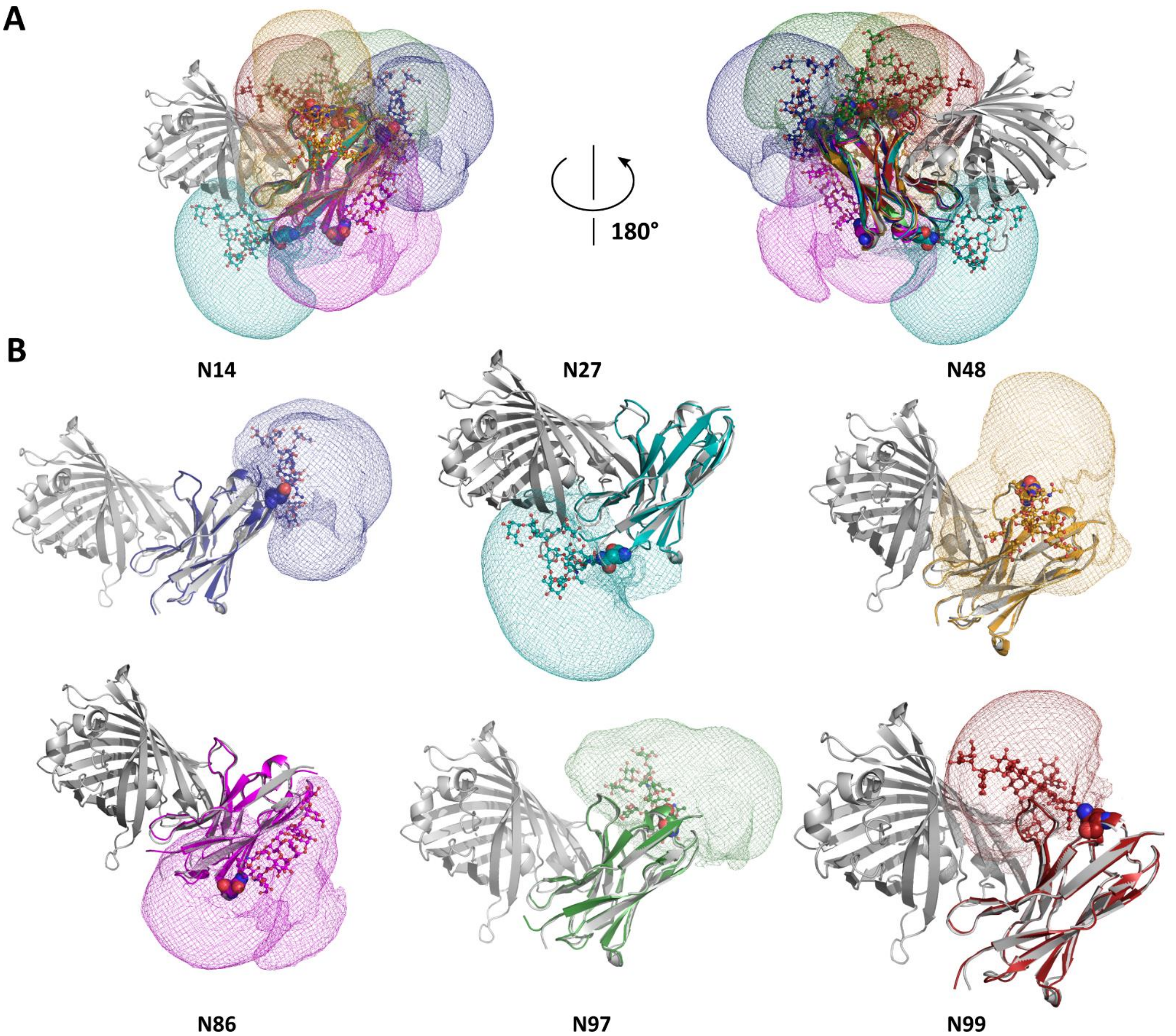


**Supplementary I. Model of each of the selected single N-glycosylation site glycovariants with a yeast Man9GlcNAc2 glycan, superimposed (A) and separately (B).** Molecular dynamics simulations of the selected GBP single N-glycosylation site glycovariants aligned to the structure of GBP with its antigen GFP (grey; PDB id 3OGO (Kubala et al., 2010)). Protein structures are represented as 'cartoon' ribbons, with glycan structures represented in ball-and-stick mode. The 0.5% space occupancy isocontours of a yeast Man9GlcNAc2 for N-glycosylation sites N14 (deep blue), N27 (teal), N48 (orange), N86 (purple), N97 (green) and N99 (red) are shown as mesh.

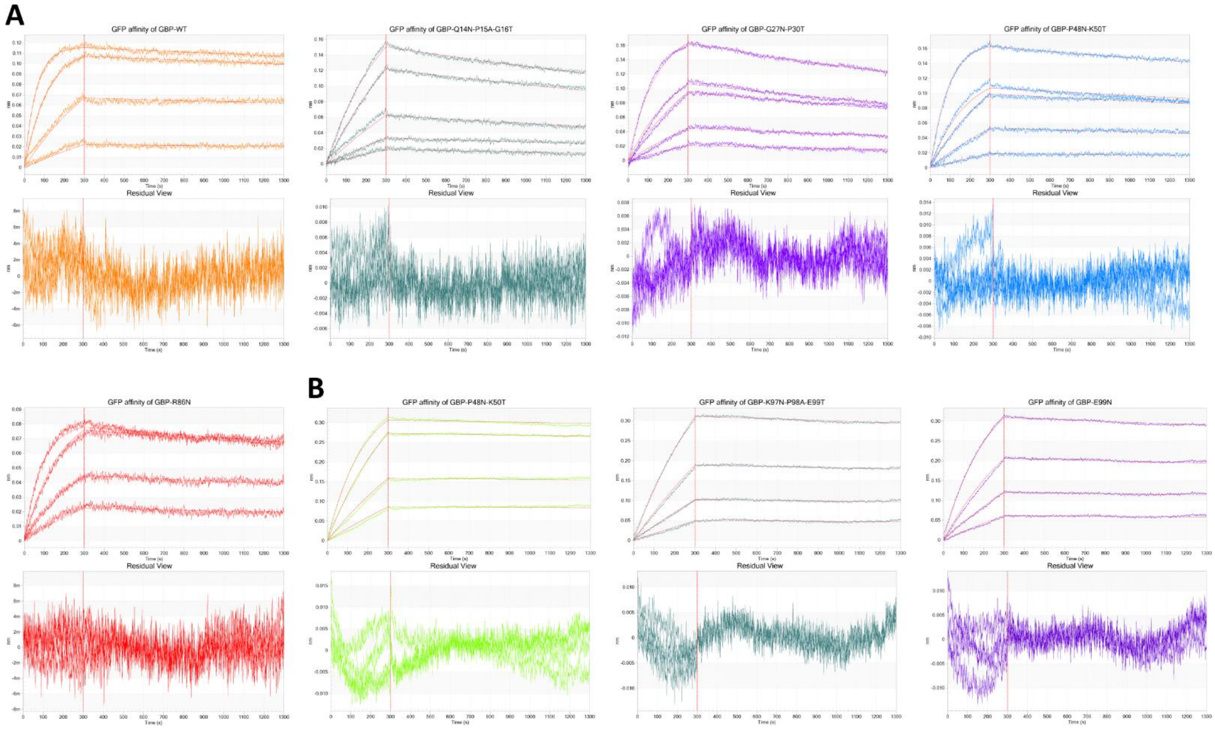


**Supplementary J. Binding of GFP is not hampered by N-glycosylation of GBP glycovariants.** Representative examples of Biolayer interferometry (BLI) curves of (A) GBP and His-6 tagged glycovariants, and (B) selected additional GBP glycovariants that carry an N-terminal EAEAGS sequence (partially processed) and C-terminal GS-His8 tag. Top curves represent the data (colored) and a 1:1 kinetics fit (red lines); bottom curves represent residuals after fit.

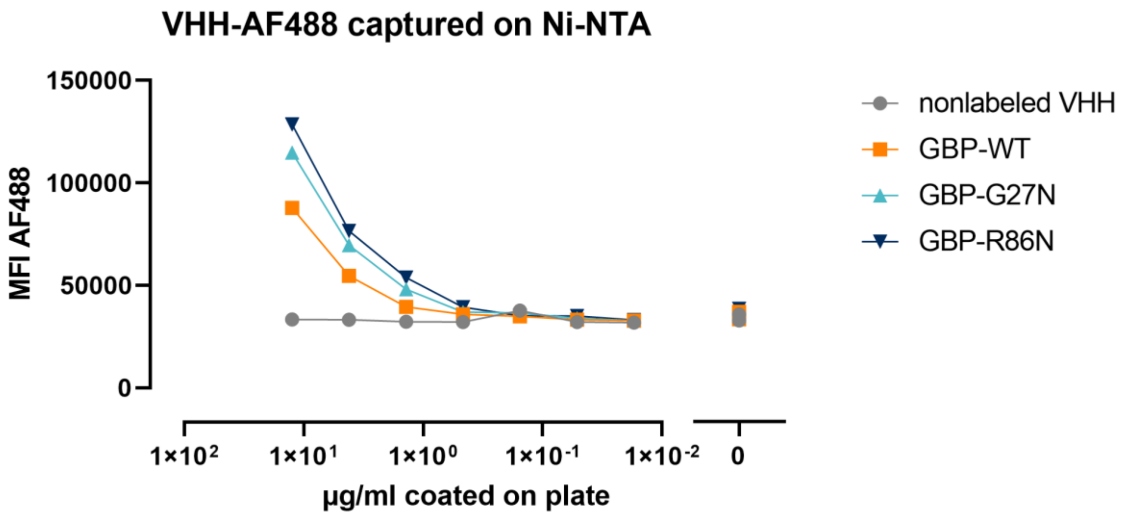


**Supplementary K. Nickel-NTA plate titration of His-tagged GBP glycovariants with fluorescence detection confirms correct AlexaFluor488 (AF488) labeling.** Three-fold dilution series of AlexaFluor488 (AF488)-labeled GBP-WT, GBP-G27N-P30T and GBP-R86N were captured on a Pierce™ Nickel Coated Plate (ThermoFisher) via their His-tag. After a PBS wash, fluorescence was detected to detect the presence of AF488-labeled His-tagged protein, using an EnVision 2105 Multimode Plate Reader (PerkinElmer).
